# Simulations predict preferred Mg^2+^ coordination in a nonenzymatic primer extension reaction center

**DOI:** 10.1101/2023.02.03.527041

**Authors:** Shriyaa Mittal, Collin Nisler, Jack W. Szostak

**Affiliations:** Howard Hughes Medical Institute, Department of Molecular Biology, and Center for Computational and Integrative Biology, Massachusetts General Hospital, Boston, MA, 02114 USA; Department of Genetics, Harvard Medical School, Boston, MA, 02115 USA; Department of Chemistry and Chemical Biology, Harvard University, Cambridge, MA, 02138 USA; Howard Hughes Medical Institute, Department of Chemistry, University of Chicago, Chicago, IL, 60637 USA

**Author notes:** Corresponding author – Jack Szostak.

## Abstract

The mechanism by which genetic information was copied prior to the evolution of ribozymes is of great interest because of its importance to the origin of life. The most effective known process for the nonenzymatic copying of an RNA template is primer extension by a two-step pathway in which 2-aminoimidazole activated nucleotides first react with each other to form an imidazolium-bridged intermediate that subsequently reacts with the primer. Reaction kinetics, structure-activity relationships, and X-ray crystallography have provided insight into the overall reaction mechanism, but many puzzles remain. In particular, high concentrations of Mg^2+^ are required for efficient primer extension, but the mechanism by which Mg^2+^ accelerates primer extension remains unknown. By analogy with the mechanism of DNA and RNA polymerases, a role for Mg^2+^ in facilitating the deprotonation of the primer 3′-hydroxyl is often assumed, but no catalytic metal ion is seen in crystal structures of the primer extension complex. To explore the potential effects of Mg^2+^ binding in the reaction center, we performed atomistic molecular dynamics simulations of a series of modeled complexes in which a Mg^2+^ ion was placed in the reaction center with inner sphere coordination to different sets of functional groups. Our simulations suggest that coordination of a Mg^2+^ ion to both O3′ of the terminal primer nucleotide and the pro-*S*_p_ non- bridging oxygen of the reactive phosphate of an imidazolium-bridged dinucleotide would help to preorganize the structure of the primer/template substrate complex to favor the primer-extension reaction. Our results suggest that the catalytic metal ion may play an important role in overcoming electrostatic repulsion between a deprotonated O3′ and the reactive phosphate of the bridged dinucleotide. Our simulations lead to testable predictions of the mode of Mg^2+^ binding that is most relevant to catalysis of primer extension.

**STATEMENT OF SIGNIFICANCE:** Prior to the evolution of complex enzymes, the replication of genetic material must have relied on nonenzymatic mechanisms. Nonenzymatic RNA template copying can be achieved through the extension of a primer by reaction with a 2-aminoimidazole (2AI) bridged dinucleotide in the presence of Mg^2+^. Despite progress in understanding the mechanism of this reaction, the catalytic role of Mg^2+^ remains poorly understood. Here, we present a series of molecular dynamics simulations of a model RNA primer-extension complex in different potential reactive conformations. We find that one configuration of both the 2AI moiety and coordination state of the Mg^2+^ promotes a geometry that is most favorable to reaction, suggesting a potential structural role for Mg^2+^ and providing insights to guide future experiments.

## INTRODUCTION

The idea that nonenzymatic RNA replication preceded the evolution of macromolecular catalysis of replication has a long history, going back to the early experimental work of Orgel et al. (1–3). Much subsequent work has focused on a search for optimal activation chemistry that would enable activated nucleotides to polymerize spontaneously on a template strand. The Orgel lab first demonstrated the template-directed polymerization of nucleoside 5′-phosphorimidazolides in 1968 (4), and then in 1982 showed that 2-methylimidazole was a superior activating group that enabled faster and more extensive copying of G/C rich templates (5). Many other potential activating groups have since been examined (6, 7). The Richert group has studied the templated polymerization of nucleotides activated by such leaving groups as N-methyl imidazole and oxyazabenzotriazole. Our laboratory identified 2-aminoimidazole (2AI) as a nucleotide activating moiety that enabled extensive copying of templates containing all four nucleotides. Furthermore, we and others have found prebiotically reasonable pathways for the synthesis of 2AI (8–11), and for the activation of nucleotides with 2AI (12–14). Given that 2AI is a potentially prebiotic activating moiety, the mechanism by which 2AI-activated nucleotides undergo template-directed polymerization is of considerable interest.

Our laboratory has extensively characterized nonenzymatic primer extension with 2AI activated monomers as a model of prebiotic template copying (15–17). Nonenzymatic RNA primer extension with 2AI activated substrates involves two chemical steps. First, two 2AI activated monomers react with each other to form an imidazolium-bridged dinucleotide (18, 19) (or a mononucleotide bridged to an oligonucleotide) which binds to the template strand adjacent to the primer. In the second step, the 3′-hydroxyl of the terminal primer nucleotide attacks the adjacent phosphate of the bridged intermediate (Figure 1A, B), leading to the formation of a new phosphodiester bond and displacing an activated nucleotide (or oligonucleotide) as the leaving group. Hence the primer is extended one nucleotide at a time whether the intermediate is an imidazolium-bridged dinucleotide or a mononucleotide bridged to an oligonucleotide (20).

**Figure 1:**
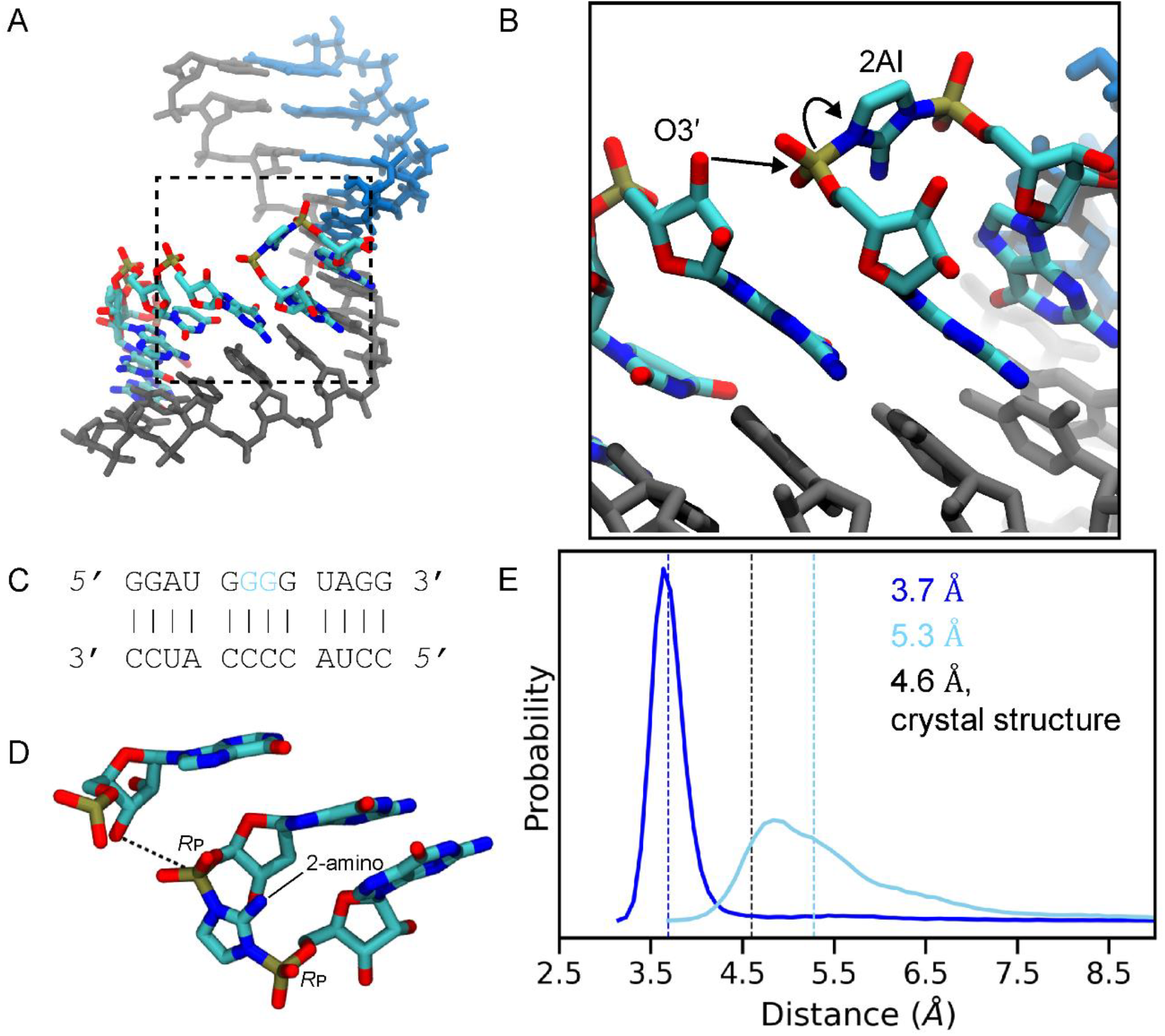
Overview of simulation system and reaction center dynamics without Mg^2+^. (A) The template is shown in gray, helper oligo in blue, and reacting imidazolium-bridged dinucleotide and primer are colored by atom type. Hydrogen atoms are not shown for clarity. (B) A zoomed in view of (A), showing the attack of the template 3′-hydroxyl on the phosphate of the dinucleotide and subsequent displacement of the activated monomer. (C) Schematic of the RNA complex used in this work. Five nucleotide long primer and helper oligonucleotides sandwich the G*G bridged dinucleotide (shown in blue) and base-pair with the 12-nucleotide template. (D) Structure of the primer extension reaction site, showing the last primer nucleotide and the G*G bridged dinucleotide. Hydrogen atoms are not shown for clarity. The dashed line shows the distance between reacting O3′ and P atoms, and *R*_p_ atoms of the backbone phosphates are indicated. (E) Probability distribution of the distance between O3′ and P atoms observed in the 3′-hydroxyl w/o Mg^2+^ (dark blue) and 3′O^-^ w/o Mg^2+^ (light blue). Dashed lines show median values for the distance distributions. The black dashed line shows the distance value of 4.6 Å in the crystal structure (21).

Kinetic studies have shown that the intermediate can form both in solution and on the template (19).

Our understanding of nonenzymatic primer extension has benefited greatly from high resolution crystal structures of RNA primer/template complexes with a template-bound imidazolium-bridged guanosine dinucleotide (G*G). In such complexes, the G*G dinucleotide is bound to the template through two canonical Watson-Crick base pairs (21). A well-ordered G*G bridged dinucleotide was observed next to the terminal primer nucleotide, with a distance of 4.6 Å between the 3′-hydroxyl and the reactive phosphate of the nucleotide in the +1 position. We were subsequently able to observe both the formation of the imidazolium-bridged intermediate from activated monomers, and its subsequent reaction with the primer, in a series of time resolved crystal structures (21). Although it has been informative to observe the two-step process of primer extension by crystallography, many aspects of the reaction mechanism remain unresolved or entirely unseen.

Perhaps the least well understood aspect of the primer extension reaction is the role of the catalytic metal ion, typically Mg^2+^. In our previous detailed kinetic studies of primer extension, the reaction does not proceed in the absence of Mg^2+^ and is initiated by the addition of Mg^2+^ at concentrations in the range of 100 mM (19). Some earlier studies have used significantly higher concentrations of Mg^2+^ to achieve optimal rates (22–24). Such high concentrations are far beyond what is needed to stabilize the primer-template duplex, or even the binding of the imidazolium bridged intermediate. Thus, it has been widely assumed that Mg^2+^ plays a critical catalytic role, but one in which binding of the catalytic Mg^2+^ ion in the reaction center is extremely weak. Presumably because of this weak binding, none of the crystal structures determined to date have captured Mg^2+^ coordinated to functional groups in the reaction center. Additionally, while crystal structures have provided a three-dimensional view into the mechanism of primer extension, crystal lattice interactions may suppress structural fluctuations that occur in solution. Structures solved to date have been determined using constructs in which critical template residues are restricted to a single sugar conformation using conformationally locked nucleotides (25, 26).

How might Mg^2+^ ions catalyze nonenzymatic primer extension? Mechanistic studies have shown that Mg^2+^ ions play vital roles in ribozyme catalysis by stabilizing leaving groups, activating nucleophiles, and coordinating non-bridging phosphate oxygens (27). In principle, Mg^2+^ ions could catalyze primer extension in similar ways. For example, inner sphere coordination with the oxygen of the 3′-hydroxyl would significantly decrease the pKa of that hydroxyl, thus favoring its deprotonation and formation of a metal-alkoxide nucleophile (28). Alternatively, Mg^2+^ coordination with a non-bridging phosphate oxygen of the bridged dinucleotide would make the phosphorus atom more electrophilic. Finally, a bridging interaction between the 3′-hydroxyl and the adjacent phosphate could decrease electrostatic repulsion between the alkoxide nucleophile and the phosphate being attacked, thereby decreasing the O3′-P distance. This is an attractive possibility since phosphodiester bond formation involves a decrease in the O3′-P distance of about 3 Å (from 4.6 Å to 1.7 Å). Understanding how this occurs will require an appreciation of the dynamic nature of the primer extension reaction site, as well as sophisticated QM simulations of the reaction pathway.

Computational studies involving molecular dynamics (MD) simulations have played a key role in understanding RNA structure, dynamics, and functions involved in processes at various lengths and timescales (29) such as base-pair opening(30), sugar puckering and bond rotations (31), dynamics of tetraloop structures (32), ion binding (33, 34), the mechanistic role of metal ions (35, 36) and riboswitch conformational changes (37). Here we study the possible mechanistic roles of Mg^2+^ in nonenzymatic primer extension. In this study, we have used MD simulations to explore the potential roles of the catalytic metal ion through its effects on the structure and dynamics of the RNA primer/template/bridged-intermediate complex. Our simulations provide an atomistic perspective of the pre-catalytic Mg^2+^-free state and of several different models of Mg^2+^-bound ground states. Our MD simulations lead to testable predictions of the metal ion coordination state that is most favorable for primer extension. Mechanistic and structural insights into how the metal ion stabilizes the catalytically competent structure may facilitate further improvements in the efficiency, extent, and accuracy of nonenzymatic primer extension.

## METHODS

A computational model of a 12 base-pair RNA duplex (5′-GGAUG G*G GUAGG-3′/5′- CCUACCCCAUCC-3′) was generated using the nab scripting language (38), where G*G represents an imidazolium-bridged guanosine dinucleotide intermediate that is bound to the -CC- template nucleotides (Figure 1A-C). We chose a G*G bridged dinucleotide and flanking guanosine residues to mimic the sequence used for crystallographic studies (21). The sequence design includes G-C terminal base pairs to stabilize the termini of the RNA duplex. The G*G bridged dinucleotide was placed within the RNA duplex by aligning the generated RNA duplex model used in our work with a previously solved crystal structure (PDB ID: 6C8E) (21). These structures were aligned using Tcl scripting in VMD (39). The five-nucleotide oligomer downstream of the bridged dinucleotide acts as a helper oligonucleotide. Downstream helper oligonucleotides have been previously suggested to contribute to primer extension by favoring a pre-organized reactive conformation of the primer-template-intermediate complex (20, 25) and also by stabilizing the RNA duplex.

In each MD simulation the RNA complex was placed in a cubic periodic box with at least 15 Å to the box boundaries and was neutralized by adding magnesium ions and a further 0.15 M MgCl_2_ in the solvent. The standard CHARMM36 (40, 41) force field was used to represent the nucleic acid and ions. The force field parameters of the imidazolium-bridged dinucleotide were developed on the basis of the CHARMM General Force Field (CGenFF) (42) through analogy to existing parameters. All oligonucleotide strands are capped with 5TER and 3TER residues within psfgen on the 5′- and 3′-end respectively, except in the case of the unprotonated 3′-end of the primer strand where after removal of the hydrogen, charge was distributed to the 3′ oxygen and neighboring atoms as previously described (43). The TIP3P water model was used to represent explicit water molecules.

The NAMD 2.14 program (44) was used to perform molecular dynamics simulations. Nonbonded interactions were smoothly switched off at 10-12 Å and long-range interactions were calculated using the Particle-Mesh Ewald (PME) method. For all simulation steps, bond distances involving hydrogen atoms were fixed using the SHAKE algorithm. Minimization was done for 10,000 steps followed by 250 ps equilibration at 300 K, followed by production MD runs. Production runs were performed in the *NpT* ensemble at 1 atm using a hybrid Nosé-Hoover Langevin piston method, and temperature was controlled using Langevin dynamics with a damping coefficient of γ = 1. The O3′-P and O2′-P distances were restrained to those observed in the crystal structure (PDB ID: 6C8E) (21)- 4.6 Å and 6.5 Å when the 2-NH_2_-Im group of the imidazolium bridge faces the major groove and 4.6 Å and 6.0 Å when the 2-NH_2_-Im group of the imidazolium bridge faces the minor groove - for the first 5 ns of the production runs. Restraints were applied using the NAMD collective variables module (45). This initial simulation stage was discarded, and the following 200 ns run was used for all analysis. Simulations were run with a 2 fs timestep, and coordinates were saved every 10 ps.

To place a Mg^2+^ ion in the reaction center, we extracted frames with the smallest O3′-P distance from our simulations without Mg^2+^ and then introduced a Mg^2+^ ion within 2 Å of the O3′-atom and either the *R*_P_ or *S*_P_ oxygen atoms on the reactive phosphate of the bridged dinucleotide using Packmol (46).

We performed five replicates for all MD systems yielding a total of 1 μs simulation time for each system. Overall, we analyzed ∼7 μs of simulation data in the current work. To prevent base-pair fraying during the simulations, we applied weak restraints on the distances d(N4,O6), d(N3,N1), and d(O2,N2) on the terminal G-C base pairs using the NAMD collective variables module (45). Canonical RNA geometries were also maintained by applying weak restraints between the C1′ atoms of the primer/template and helper/template nucleotides two nucleotides upstream and downstream of the nicked backbone.

The structural analysis of the MD trajectories was performed using CPPTraj V5.1.0 (47), MDTraj 1.9.4 (48), and VMD (39). Root mean-squared deviation (RMSD) was calculated by superposing the simulation frame onto the first frame of the simulation trajectories prior to minimization. RMSD values were calculated for all nucleic acid atoms and bridged dinucleotide atoms (except hydrogen atoms). Sugar puckering was calculated from the five torsion angles of the five-membered sugar ring. The torsion angles were converted to the pseudorotation phase angle and amplitude parameters based on the Altona-Sundaralingam definition (49). Each point on the circular histogram plots corresponds to a value of the phase angle (0 - 360°) moving clockwise from a vertical value of 0° and the amplitude of the pucker (0 - 70°) which increases radially from the center of the plot. Ribose rings avoid a planar conformation and are usually present in the C2′-endo or C3′-endo conformation.

The CHARMM force field has been considered by some to be less accurate than the AMBER force field for MD simulations of nucleic acids (29). We therefore carried out control simulations on the same major groove-facing-2AI configuration as described above, starting from the same initial coordinates, but using the AMBER OL3 force field (50) to simulate the RNA duplex primer, template, and helper nucleotides. Parameters were converted to AMBER using the CHARMM-GUI Force Field Converter (51, 52), and simulations were run using NAMD 2.14 using the TIP3P explicit model of water and 12-6-4 ion force field. Bonded parameters for the imidazolium-bridged dinucleotide, including bonds, angles, and dihedrals, were taken from those generated in CGenFF. Nonbonded parameters for the imidazolium group were generated using OpenFF, while OL3 parameters were used to represent the remaining non-bonded terms for the bases of the imidazolium-bridged dinucleotide. The final system was minimized for 10,000 steps, followed by 250 ps of equilibration in which the nucleic acid backbone heavy atoms were constrained with a force constant of 1 kcal mol^-1^ Å^-2^ and non-backbone nucleotide heavy atoms were constrained with a force constant of 0.5 kcal mol^-1^ Å^-2^. Production MD was performed in the *NpT* ensemble at 1 atm using a hybrid Nosé-Hoover Langevin piston method. Simulation parameters, including temperature and pressure control, PME, hydrogen atom constraints, and timestep were implemented exactly as above, except for a 1-4 scaling term of 0.833333, a cutoff of 9 Å, and rigid Tolerance of 0.0005, as suggested by NAMD developers when using the AMBER force field (ks.uiuc.edu). Analysis of the control simulation was performed as described above.

Finally, the lack of polarization or charge-transfer effects in standard force fields can result in inaccurate thermodynamic and kinetic properties of Mg^2+^ binding in aqueous systems (29, 53). To better characterize the RNA backbone-Mg^2+^ interaction in our simulations and to validate the classical simulations, we carried out mixed quantum-mechanical/molecular mechanical (QM/MM) simulations on the same major groove-facing-2AI with *S*_P_ and deprotonated O3’ oxygen bound Mg^2+^ configuration as described above using NAMD’s QM/MM interface (54).

The 3’ primer sugar, 2AI-bridged backbone up to the 5’ carbons of each monomer in the bridged dimer, the interacting Mg^2+^, and 4 inner-sphere coordinating water molecules were all treated at the QM level (49 atoms total; Figure S13A), resulting in a multiplicity of 1 and no net charge for the QM region. The rest of the system was treated at the classical level using the CHARMM force field as described above. The QM calculations were performed by ORCA (55) using the HF-3c semi-empirical Hartree Fock method (56). Interactions between the QM and MM region were treated with an electrostatic embedding scheme, and covalent bonds split at the QM/MM boundary were treated with a Charge Shifting method. Charge distribution in the QM region was taken from ORCA and updated at every step using the mulliken charge calculation mode. Other than the use of a 0.5 fs timestep, parameters for the MM region were identical to those described above for CHARMM simulations. Analysis of the QM/MM simulation was performed as described above. Self-consistent field (SCF) electron density and electrostatic potential maps were generated using the orca_plot command and the molecular orbitals output by ORCA, and plotted with an isovalue of 0.085.

## RESULTS AND DISCUSSION

### Effect of protonation state of the primer 3′ hydroxyl in the absence of Mg^2+^

Previous work has suggested that the active nucleophilic species in nonenzymatic primer extension is a 3′-alkoxide. This conclusion was supported in part by the absence of any significant solvent deuterium isotope effect, which is consistent with the necessary proton transfer occurring prior to the transition state. In addition, the pH dependence of the reaction rate has been interpreted as reflecting the metal-ion dependent deprotonation of the 3′-hydroxyl (28). However, these experiments are indirect, and the possibility that the nucleophile is the protonated hydroxyl cannot be fully discounted. We have therefore modeled the ground state of the reaction center with the 3′-hydroxyl either protonated or deprotonated. In principle the 3′- hydroxyl could be deprotonated by proton transfer to a Mg^2+^ coordinated to 5 water molecules and a hydroxide ion, such that O3′ would not be inner-sphere coordinated to the Mg^2+^ ion. Such an alkoxide would be more nucleophilic than a directly metal coordinated alkoxide, and much more nucleophilic than a protonated 3′-OH (27). To provide atomistic insight into the dynamics of the ground state and structural implications of a deprotonated O3′, we first compared MD simulations in which O3′ was a protonated 3′-OH vs. an O3′ alkoxide, both in the absence of Mg^2+^.

We first carried out simulations with the conformation of the bridged dinucleotide such that the 2-amino group of the 2-aminoimidazolium was hydrogen bonded with the *R*_P_ oxygen atoms of the flanking phosphates (2-NH_2_-Im:*R*_P_, Figure 1D), referred to as the major groove-facing conformation in Zhang et al. (21). Overall, the structure in simulations when O3′ is protonated show low root-mean-square deviation (RMSD) with respect to the starting conformation indicating excellent maintenance of the double helical structure (Figure S1A, B). The higher RMSD of the reaction center when O3′ is deprotonated indicates that the deprotonated species makes the reaction center slightly more dynamic (Figure S1C, D).

To further characterize the reaction center and understand the origin of the increased flexibility of the reaction center in the deprotonated case, we compared the distance between the O3′ atom of the terminal primer nucleotide and the P of the adjacent phosphate of the bridged dinucleotide (Figure 1D). Shorter distances, which would favor the primer extension reaction, are observed when O3′ is protonated, while the median distance increases from 3.7 Å to 5.3 Å in the deprotonated case (Figure 1E). In our most recent crystal structure (PDB: 6C8E (21)), the corresponding distance is 4.6 Å. Crystal structures solved with G(5’)ppp(5’)G – an analog of the imidazolium bridged dinucleotide – in the absence of helper oligos (PDB: 5UED (57)) show the distance between O3′ and the nearest phosphorous atom to be 4.1 Å. In the presence of helper oligos (PDB: 6AZ4 (25)) the corresponding distance is significantly shorter, 3.7 Å. Our protonated O3′-P distances lie within the range of those observed in these crystal structures, but this key distance increases significantly when the 3′-hydroxyl is deprotonated.

To examine the effect of the protonation state of the 3′-hydroxyl on the ribose conformation, we calculated the pseudorotation angles of the ribose sugar of the terminal primer nucleotide. Figure S2 shows the simulated conformational ensemble projected on a circular plot where the phase angle increases clockwise from a vertical value of 0° and the puckering amplitude increases radially from the center. The resulting probability density plot shows that the most prominent conformation observed is the C3′-endo for both protonated and deprotonated O3′ simulations.

During our MD runs, the ribose sugar exhibits reversible conformational switches to the C2′- endo conformation (Figure 2C, D). As seen in Figure S2, this conformational change from C3′- endo to C2′-endo occurs through the O4′-endo conformation. The C2′-endo conformation is observed for longer periods in the deprotonated case and the time spent in this conformation increases from 3.5% when O3′ is protonated to 30% when O3′ is deprotonated. In contrast, the terminal primer nucleotide sugars in the crystal structure (PDB: 6C8E (21)) are all in the C3′- endo conformation (Figure S2A).

**Figure 2:**
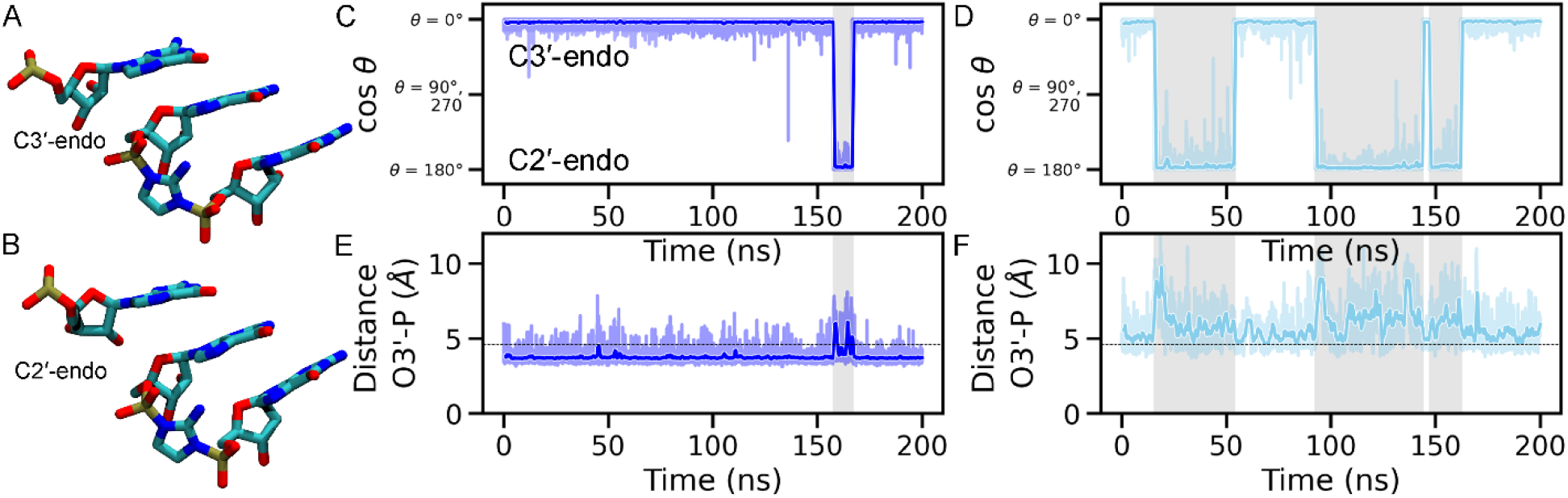
Correlation of sugar conformation and O3′-P distance during MD simulations without Mg^2+^. (A,B) Molecular figures showing the conformation of the terminal primer sugar in the (A) C3′-endo conformation and (B) C2′-endo conformation. (C, D) Time series of the primer 3′-nucleotide sugar pucker pseudorotation angle for the (C) 3′-hydroxyl and (D) 3′O^-^ simulation systems, without Mg^2+^ in the reaction center. Sugar pucker angles close to 0° correspond to the C3′-endo sugar conformation and angles close to 180° correspond to the C2′- endo sugar conformation. Time series for cosine of the angle (cos θ) are shown for clarity and angle (θ) values are indicated. (E,F) Time series of the distance between O3′ (primer) and P (bridged dinucleotide) for the (E) 3′-hydroxyl and (F) 3′O^-^ simulation systems, both without Mg^2+^ in the reaction center. The horizontal dashed lines indicate the crystal structure value. Intervals with larger distance values and C2′-endo sugar conformations are highlighted in gray. For all time series plots, dark traces show the data averaged over a 1 ns window, while the lighter envelope shows the full range of the data recorded at 10 ps time steps in our simulations. All simulation replicates are shown in Figure S3 and S4.

During the simulation runs, periods of high O3′-P distance are highly correlated with the primer ribose conformation switch to C2′-endo (Figures 2, S3, and S4), in both protonated and deprotonated cases. It is unclear whether the increase in distance is a cause or consequence of the sugar conformation switch. Nevertheless, the correlation explains why modified primer 3′- nucleotides with sugars with an increased preference for the C3′-endo conformation are better at primer extension (26, 58, 59) because this conformation is associated with shorter distances of attack. The larger O3′-P distances and the prevalence of the C2′-endo sugar conformation observed when O3′ is deprotonated in our simulations suggest that deprotonation without a bound Mg^2+^ (while highly unlikely to occur) would be anti-catalytic. This observation is consistent with the possibility that the deprotonated O3′ primer is stabilized by coordination with a metal ion, which may act to overcome the electrostatic repulsion between the deprotonated O3′ and the phosphate it is attacking.

### Orientation of the 2-aminoimidazole affects the O3′-P distance and angle of attack

To examine the effect of the orientation of the imidazolium moiety of the bridged dinucleotide on the geometry of the reaction center, we performed MD simulations beginning with the minor groove-facing conformation, in which the imidazolium 2-NH_2_-Im group forms hydrogen bonds with the *S*_P_ oxygens of the flanking phosphates (2-NH_2_-Im:*S*_P_, Figure 3A). As before, we performed these simulations for both the protonated and deprotonated states of the primer 3′ hydroxyl. As in the major groove-facing orientation discussed above, the minor groove-facing orientation simulations show a smaller O3′-P distance of attack when O3′ is protonated (Figure 3B). When O3′ is deprotonated, the median distance increases by 1.7 Å, similar to the 1.6 Å increase seen in the major groove-facing simulations discussed previously.

**Figure 3:**
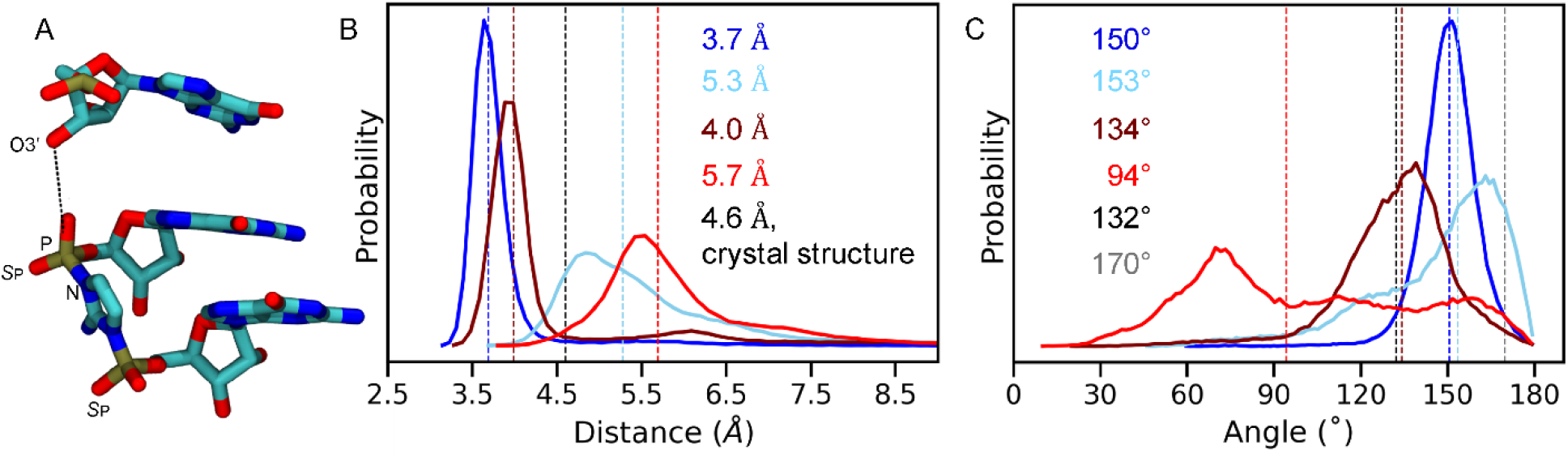
O3′-P distance and angle of attack during MD simulations in different 2AI conformations without Mg^2+^. (A) Structure of the primer extension reaction site, showing the last primer nucleotide and the G*G bridged dinucleotide with the imidazolium 2-NH_2_-Im pointing towards the minor groove. Hydrogen atoms are not shown for clarity. The dashed line shows the O3′-P distance. (B) Probability distribution of the distance between O3′ and P atoms observed in the four simulation ensembles: major groove-facing 2-NH_2_-Im conformations for 3′- hydroxyl w/o Mg^2+^ (dark blue) and 3′O^-^ w/o Mg^2+^ (light blue) and minor groove-facing conformations for 3′-hydroxyl w/o Mg^2+^ (brown) and 3′O^-^ w/o Mg^2+^ (red). Colored dashed lines show median values for the distance distributions. The black dashed line shows the crystal structure distance. (C) Probability distribution of the angle of attack, measured using the O3′-P-N atoms, in the four simulation ensembles. Dashed lines show median values for the distributions. The black and gray dashed lines show the crystal structure angle for the major groove and minor groove-facing conformations, respectively.

The minor groove-facing orientation of the imidazolium group has relatively minor effects on the ribose pucker of the 3′-primer nucleotide. In both protonation states, the sugar is predominantly in the C3′-endo conformation (Figure S5). However, when O3′ is deprotonated, the terminal primer sugar shows a slightly higher probability of a C3′-endo conformation as opposed to when O3′ is protonated, which is opposite to the trends observed for the major groove-facing imidazolium state (comparing Figures S2 and S5). The sugar conformation is also not as well correlated with the O3′-P distance. We suggest that the increased distance between the primer and the bridged dinucleotide causes the sugar pucker and the O3′-P distance to vary independently of each other. The first sugar of the bridged dinucleotide (G1) is predominantly C3′-endo for the simulations with a major groove-facing imidazolium, but predominantly C2′- endo for the minor-grove facing imidazolium orientation (Figure S6A). In contrast the ribose of the second sugar (G2) is always predominantly C3′-endo (Figure S6B). Both G1 and G2 sugars in crystal structures with the G*G bridged dinucleotide and GpppG are in the C3′-endo conformation. Overall, in the four simulation systems discussed so far, the maximum probability for the C3′-endo sugar conformation for all three ribose sugars is seen when the primer O3′ is protonated and the bridged dinucleotide is in the major groove-facing orientation.

The orientation of the imidazolium moiety also affects the angle of attack between the 3′- hydroxyl and the P-N bond to be broken during phosphodiester bond formation. A more favorable angle for in-line attack (closer to 180°) is primarily seen in the major groove-facing conformations (Figure 3C). This is in contrast with the crystal structures, where the angles of attack are 132° and 170° for the major groove and minor groove-facing conformations, respectively. While the cause of this discrepancy is unclear, it may reflect an artifact of fitting both possible orientations into the same density in the crystal structures, while MD simulations provide insights into the relaxed conformations explored during the time course of the simulations.

The O3′-P distance, the terminal primer sugar pucker, and the angle of attack are key structural characteristics of the primer extension reaction center. In summary, we observe a shorter O3′-P distance, increased C3′-endo conformation of the terminal primer sugar, and a higher than ∼150° angle of attack when the imidazolium 2-NH_2_-Im faces the major groove of the RNA duplex. All three parameters shift towards less catalytically favorable values when the 2-NH_2_-Im faces the minor groove. Therefore, while both conformations are likely to occur, we suggest that the major groove-facing conformation of the bridged dinucleotide imidazolium is more favorable for phosphodiester bond formation.

Our MD simulations suggest that the major groove-facing conformation of the bridging 2- aminoimidazolium group may be stabilized by water-mediated coordination with a Mg^2+^ ion. In a crystal structure (PDB ID: 6C8D (21)) in which two template-bound 2′-deoxyguanosine-5′- monophosphate monomers sit next to the primer in an RNA duplex, a metal ion (Sr^2+^ ion in PDB ID: 6CAB (21)) forms water-mediated contacts with N7 and O6 of the 3′-terminal primer guanosine and with the N7 atoms of the G1 and G2 template-bound monomers. Both inner-sphere and outer-sphere interactions of Mg^2+^ with the N7 atom of guanosine have been observed in a large number of solved RNA crystal structures (60). Interestingly, in our MD simulations, we observe water-mediated interactions between a Mg^2+^ ion and N7 of the first guanosine (G1) of the bridged dinucleotide, and with its O6 atom. Additionally, the same metal ion forms water-mediated interactions with the primer 3′ guanosine and the G2 nucleobase of the bridged dinucleotide which supports its previously suggested role in the better copying chemistry of G*G as opposed to other nucleobases. Moreover, it is possible that this Mg^2+^ may also form a water-mediated contact with the 2-amino group of the bridging imidazolium when it is in the major groove-facing conformation (Figure S7A). This imidazolium-ion contact is mediated through water molecules and does not occur when the imidazolium faces the minor groove (Figure S7B). Hence, we posit that a secondary metal ion may help to pre-organize the catalytically favorable major groove-facing conformation of the bridged dinucleotide.

To determine the extent to which the observed dynamics are dependent on the choice of force field, a 200 ns equilibration simulation without constraints was performed on the major groove-facing system with protonated O3′ using the AMBER force field (see methods for simulation details). For analysis, the simulation was split into two 100 ns sub-trajectories to test for self-consistency (61). Compared to the first 100 ns of the first replicate simulated using the CHARMM force field on the same major groove-facing configuration, the AMBER system exhibited similar terminal primer sugar puckering (Figure S8A-C), RMSD of nucleic acid heavy atoms (Figure S8D), O3′-P distance (Figure S8E-F), O-P-N attack angle (Figure S8G), and stability of WC base pairing in sites flanking the imidazolium-bridged dinucleotide (Figure S8H). While some variability is observed in the degree of fluctuation between systems, the probability distributions show similar mean values (Figure S8D-F, insets). Additionally, similar variability is observed between sub-trajectories using the CHARMM force field (Figure S3, first and second rows). Finally, the terminal primer sugar in the CHARMM simulation does exhibit a greater propensity for switching to a C2′-endo conformation compared to the AMBER simulation. However, this simulation replicate exhibited the highest conformational dynamics in the terminal primer sugar of all the CHARMM simulation replicates (Figure S3), suggesting the sugar conformational ensemble seen in the AMBER simulation is comparable to the CHARMM simulations in aggregate (Figure S2A). Overall, these results suggest that the dynamic effects presented in previous and subsequent simulations are the result of differences in the systems being investigated and not the simulation methods used.

### Effects of Mg^2+^ coordination to the primer O3′ and the adjacent phosphate of the bridged dinucleotide

To identify potentially important interactions of a Mg^2+^ ion in the reaction center, we introduced a Mg^2+^ ion with inner-sphere interactions with a deprotonated O3′ and either the *S*_P_ or *R*_P_ oxygen atoms on the reactive phosphate of the bridged dinucleotide. These systems will be referred to as 3′O^-^ Mg^2+^@*S*_P_, 2-NH_2_-Im:*R*_P_ (Figure 4A) and 3′O^-^ Mg^2+^@*R*_P_, 2-NH_2_-Im:*R*_P_ (Figure 4B), respectively. We also performed simulations with the Mg^2+^ coordinated to the *R*_P_ oxygen and the 2-NH_2_-Im facing the minor groove (3′O^-^ Mg^2+^@*R*_P_, 2-NH_2_-Im:*S*_P_) (Figure 4C). When we attempted to start our MD simulations in the remaining 3′O^-^ Mg^2+^@*S*_P_, 2-NH_2_-Im:*S*_P_ arrangement, the Mg^2+^ ion spontaneously switched its coordination from the *S*_P_ to the *R*_P_ oxygen. In all replicates of our three Mg^2+^ bound simulations the ion forms stable bridging interactions between the terminal primer nucleotide and the bridged dinucleotide.

**Figure 4:**
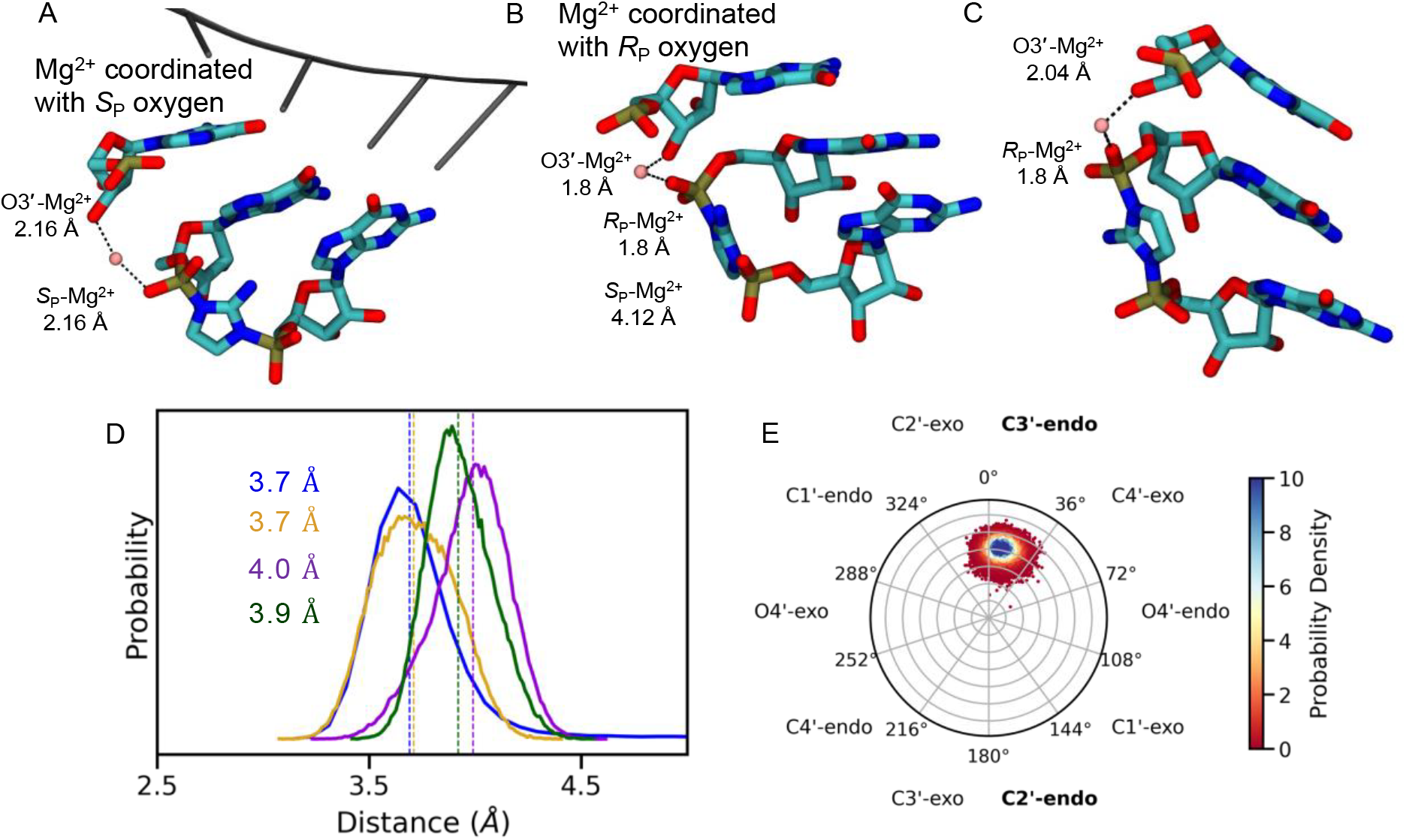
Representative snapshots showing plausible Mg^2+^ coordination geometry and their effects on dynamics. (A) 3′O^-^ w/ Mg^2+^@*S*_P_, 2-NH_2_-Im:*R*_P_ with the template residues visualized in gray, (B) 3′O^-^ w/ Mg^2+^@*R*_P_, 2-NH_2_-Im:*R*_P_, and (C) 3′O^-^ w/ Mg^2+^@*R*_P_, 2-NH_2_- Im:*S*_P_. (D) Probability distribution of the distance between O3′ and P atoms observed in the three simulation ensembles with Mg^2+^ in the reaction center. The simulation ensembles are 3′O^-^ w/ Mg^2+^@*S*_P_, 2-NH_2_-Im:*R*_P_ (yellow), 3′O^-^ w/ Mg^2+^@*R*_P_, 2-NH_2_-Im:*R*_P_ (purple), and 3′O^-^ w/ Mg^2+^@*R*_P_, 2-NH_2_-Im:*S*_P_ (green). Distance probability distribution for 3′-hydroxyl w/o Mg^2+^, 2- NH_2_-Im:*R*_P_ (dark blue) is shown for comparison. Colored dashed lines show median values for the distance distributions. (E) Circular histogram of pseudorotation angles of the terminal primer nucleotide sugar for the 3′O^-^ w/ Mg^2+^@*S*_P_, 2-NH_2_-Im:*R*_P_ simulation case. All Mg^2+^ coordination geometries with respect to the oxygen atoms are comparable and Figure S17 shows that the O3′- Mg^2+^-O angles are close to 95° for all Mg^2+^ coordination geometries.

Comparison of the 3′O^-^ Mg^2+^@*S*_P_, 2-NH_2_-Im:*R*_P_ simulation (yellow curve in Figure 4D) with the corresponding protonated simulation ensemble lacking a bound Mg^2+^ ion (dark blue curve in Figure 4D) shows a similar unimodal O3′-P distance distribution with the same median of 3.7 Å. The median O3′-P distance of 4.0 Å is slightly larger for the 3′O^-^ Mg^2+^@*R*_P_, 2-NH_2_-Im:*R*_P_ simulation (purple curve in Figure 4D). The 0.3 Å increase in the median O3′-P distance is too small to indicate that Mg^2+^ coordination with one or the other of the phosphate oxygen atoms would significantly favor the primer extension reaction. However, in the latter case where both the 2-NH_2_-Im and the Mg^2+^ are coordinated to the same oxygen atom, the reaction might be affected for other reasons, such as effects on the nucleophilicity of the 3′O^-^ or the electrophilicity of the reactive phosphorous atom.

Our previous simulations in the absence of bound Mg^2+^ showed that deprotonation of the 3′- hydroxyl is correlated with longer O3′-P distances. However, when there is a Mg^2+^ ion bridging the primer and the phosphate, the O3′-P distances in our MD simulations greatly decrease, with the median distance decreasing from 5.7 Å (red curve in Figure 3B) to 3.9 Å (green curve in Figure 4D). Hence, although deprotonation was anti-catalytic in the absence of Mg^2+^, a bridging Mg^2+^ atom coordinating with the deprotonated O3′ overcomes this effect and plays an important role in preventing the primer and bridged dinucleotide from moving apart from each other. This effect of Mg^2+^ is independent of which of the oxygen atoms of the bridged dinucleotide the Mg^2+^ coordinates with, or the orientation of the imidazolium 2-NH_2_-Im.

To further characterize the ground state of the primer/template/bridged-intermediate complex we have also examined the terminal primer sugar conformation, angle of attack, and RMSD of the reaction center in the presence of Mg^2+^. In addition to decreasing the O3′-P distance, the presence of a bridging Mg^2+^ ion is correlated with the terminal primer nucleotide sugar being almost exclusively in the C3′-endo conformation (Figures 4E and S9), which has been shown experimentally to favor primer extension (58, 59). In simulations where the imidazolium 2-NH_2_- Im group faces the major groove the G1 sugar pucker is predominantly C3′-endo, but when the imidazolium 2-NH_2_-Im group faces the minor groove, the G1 nucleotide sugar pucker conformation shifts toward being C2′-endo (Figure S10A), consistent with our simulations without Mg^2+^ in the reaction center (Figure S6A). In the simulation in which both the Mg^2+^ ion and the imidazolium 2-NH_2_, are interacting with the *R*_P_ oxygen of the phosphate, the G1 sugar adopts a range of conformations including C4′-exo, C1′-exo, and C2′-endo, with very low probability C3′-endo sugar puckering (Figure S10A). As seen in simulations without Mg^2+^ the sugar pucker of the G2 nucleotide does occasionally adopt the C2′-endo conformation but the C3′-endo conformation shows a larger population density (Figure S10B). Recent work has shown that the primer extension reaction is faster when the bridged dinucleotide sugars are in the C3′-endo conformation (62). From our simulations, only the 3′O^-^ w/ Mg^2+^@*S*_P_, 2-NH_2_-Im:*R*_P_ system exhibits a primarily C3′-endo conformation for both G1 and G2.

The 3′O^-^ w/ Mg^2+^@*S*_P_, 2-NH_2_-Im:*R*_P_ system also shows a favorable angle for in-line attack (Figure 5A; median value of 148°, yellow curve in Figure 5B) which is comparable to the median value of 150° for the simulations with protonated O3′ without a Mg^2+^ ion (dark blue curve in Figure 5B). All other simulation systems with Mg^2+^ show less favorable angles of attack. Overall, a reaction center organization in which the imidazolium group faces the major groove and the Mg^2+^ ion forms inner-sphere contacts with the *S*_P_ oxygen of the reactive phosphate would appear to favor the primer extension reaction. Moreover, a lower RMSD of the base-paired primer/template and helper/template regions (Figure S11A) and the bridged-dinucleotide (Figure S11B) region for the 3′O^-^ w/ Mg^2+^@*S*_P_, 2-NH_2_-Im:*R*_P_ simulations suggest that the presence of the bridging metal ion in the reaction center has a global effect in stabilizing the primer extension complex.

**Figure 5:**
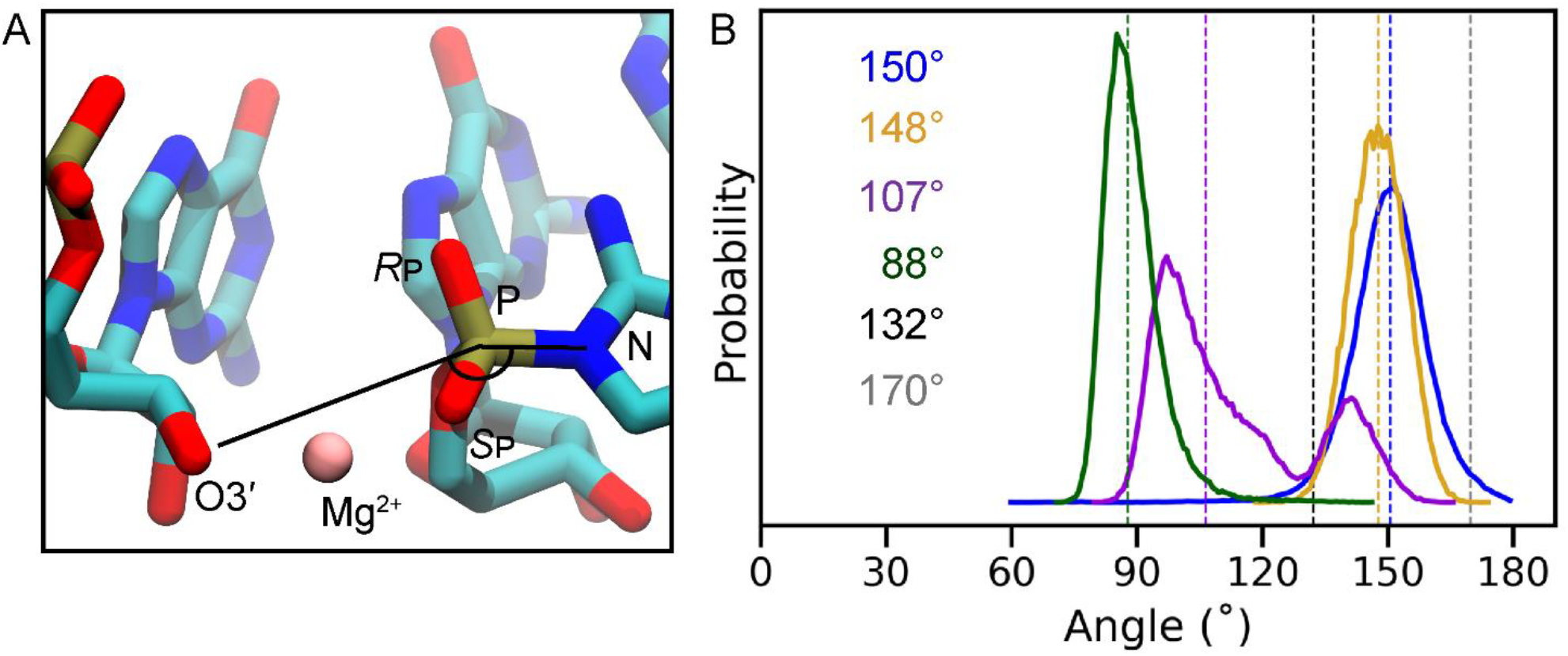
Effect of 2AI conformation and Mg^2+^ binding sites on angle of attack. (A) Molecular figure of the 3′O^-^ w/ Mg^2+^@*S*_P_, 2-NH_2_-Im:*R*_P_ system showing the angle of attack measured throughout the simulations. (B) Probability distribution of the angle of attack, measured among O3′-P-N atoms in four simulation ensembles: 3′-hydroxyl w/o Mg^2+^, 2-NH_2_- Im:*R*_P_ (dark blue), 3′O^-^ w/ Mg^2+^@*S*_P_, 2-NH_2_-Im:*R*_P_ (yellow), 3′O^-^ w/ Mg^2+^@*R*_P_, 2-NH_2_-Im:*R*_P_ (purple) and 3′O^-^ w/ Mg^2+^@*R*_P_, 2-NH_2_-Im:*S*_P_ (green). Dashed lines show median values for the distributions. The black and gray dashed line shows the crystal structure angle for the major groove and minor groove-facing conformations, respectively.

Interestingly, we also observed that when O3′ is deprotonated and no Mg^2+^ is present in the reaction center, the average O2′-P distance is less than the average O3′-P distance, which might suggest an increased probability of forming a 2′-5′ linkage. Previous studies from our laboratory have shown that primer extension by reaction with an imidazolium bridged dinucleotide intermediate increases the proportion of 3′-5′ linkages formed, compared to reaction with an activated monomer (63). Our results suggest that through its role in stabilizing the conformation of the bridged dinucleotide, the metal ion also prevents the formation of the incorrect 2′-5′ extended product. Furthermore, the predicted favorable metal coordination with the *S*_P_ oxygen atom of the bridged dinucleotide (yellow symbols in Figure S12) typically shows a more significant difference between the average O3′-P and O2′-P distances, increasing the likelihood of formation of the 3′-5′ linkage.

To further validate models used for the interaction between RNA and Mg^2+^, a 100 ps QM/MM simulation was performed on the 3′O^-^ Mg^2+^@*S*_P_, 2-NH_2_-Im:*R*_P_ system, with the reaction center modeled at the HF-3c semi-empirical Hartree-Fock level (Figure S13A; see methods for simulation details). The QM/MM simulation exhibited similar O-P-N angle (Figure S13B) compared to the CHARMM simulations (Figure 5B), and average O3’-P distance was 3.8 Å in the QM/MM simulation (Figure S13C) compared to 3.7 Å in the CHARMM simulation (Figure 4D), suggesting these parameters are well modeled without the effects of polarizability. The O3’- Mg^2+^-O(*S*_P_) angle was measured (Figure S13D and E), and was similar to the angle observed in the CHARMM simulations between the same 3 atoms (Figure S17). Finally, the QM/MM system exhibited similar terminal primer sugar puckering (Figure S13F) compared to the CHARMM simulations (Fig. S9A), though the QM/MM simulation exhibited a slight preference for a C4’- exo conformation relative to the CHARMM simulation. While the timescale limitations make it difficult to discern whether this is a favorable conformation, or the system is in transition from one state to another, experiments and simulations have shown the C4’-exo conformation to be significantly populated in other RNA systems (64). A potential molecular explanation for this deviation can be found in the polarizability of the QM/MM system (Figure S13G and H), in which the calculated charge of the H2’ atom was 0.33 at the end of the QM/MM simulation, while the same atom had a fixed charge of 0.09 in the MM simulation. This resulted in a tighter interaction between the H2’ and O3’ in the QM/MM simulation (Figure S13I), which biases the geometry of the sugar towards a more planar configuration. Despite this difference in charge between the classical and QM/MM simulations, classical modeling of essential geometric parameters such as O3’-P distance, O-P-N angle, and O3’-Mg^2+^-O(*S*_P_) angle agree with the quantum calculations over the timescale sampled.

The structural features expected for optimal reaction center geometry such as short O3′-P distance, C3′-endo sugar conformations, in-line angle of attack, and 3′-5′ regioselectivity are either absent or not present for extended timescales in the 3′O^-^ w/ Mg^2+^@*R*_P_, 2-NH_2_-Im:*S*_P_ and 3′O^-^ w/ Mg^2+^@*R*_P_, 2-NH_2_-Im:*R*_P_ simulation systems. In the latter case, the hydrogen bond between the 2-NH_2_-Im and the *R*_P_ oxygen is likely to be weakened due to the Mg^2+^ being coordinated to the same oxygen atom (Figure S14E). Thus Mg^2+^ coordination with the *S*_P_ oxygen and 2-NH_2_-Im hydrogen bonded to the *R*_P_ oxygen– the major groove-facing orientation - is likely to be the preferred geometry for the preorganization of the reaction center.

## CONCLUSION

Understanding the structure and dynamics of the primer extension reaction center may give insight into how to improve the rate (18) and fidelity (65) of primer extension. In the present study, we performed MD simulations of a primer/template/bridged dinucleotide complex, in the absence or presence of a catalytic Mg^2+^ in the reaction center. We modeled the system with both a protonated and a deprotonated 3′-hydroxyl, and with both likely orientations of the imidazolium moiety of the bridged dinucleotide intermediate. By analogy with the mechanism of enzyme catalyzed primer extension, one possible model for nonenzymatic primer extension is that inner-sphere coordination of the 3′-hydroxyl with Mg^2+^ lowers the pKa of the hydroxyl, thereby facilitating its deprotonation to form an alkoxide nucleophile. Our MD simulations are consistent with a role for a bridging Mg^2+^ in overcoming the electrostatic repulsion between the ionized O3′^-^ and the negatively charged phosphate. In our simulations, the median distance from the primer O3′^-^ to the phosphorous atom of the adjacent phosphate is 3.7 Å in the presence of Mg^2+^ in the reaction center, but much greater in the absence of a coordinated Mg^2+^ ion. Along with the organizing role of Mg^2+^ in the reaction center, the major groove-facing conformation of the imidazolium moiety of the bridged dinucleotide is correlated with an increased fraction of the C3′-endo sugar conformation and a higher in-line angle of attack. Our comparison of Mg^2+^ coordination with either of the two non-bridging phosphate oxygen atoms, *S*_P_ and *R*_P_, of the bridged dinucleotide suggests that an inner-sphere coordination with the *S*_P_ oxygen atom is structurally favorable for the catalytic step. We are currently testing our predicted 3′O^-^ w/ Mg^2+^@*S*_P_ coordination by phosphorothioate metal rescue experiments (36, 66) and X-ray crystallography.

Based on studies of primer extension with modified 3′-terminal sugars, Giurgiu et al. proposed that the primer extension reaction involves an increase in the puckering amplitude of the terminal sugar as the O3′ atom attacks the adjacent phosphorous atom of the bridged dinucleotide (26). In some of our MD simulations we do see an association of larger puckering amplitude with configurations with a shorter O3′-P distance. Interestingly this trend is strongest in the system proposed above to be most favorable for primer extension, i.e. the 3′O^-^ w/ Mg^2+^@*S*_P_, 2-NH_2_- Im:*R*_P_ system, where Mg^2+^ is coordinated to the *S*_p_ oxygen and the amine of the bridging 2- aminoimidazolium is hydrogen bonded with the *R*_p_ phosphate oxygen (Figure S15 and S16A).

Since modified sugars can significantly impact the rate of the primer extension reaction, future computational studies incorporating sugars such as ANA, DNA, and LNA in the primer/template/bridged dinucleotide complex may reveal the contribution, if any, of increasing sugar pucker amplitude and related structural factors to the rate of the primer extension reaction (26, 67–69). Because transient intermediate conformations cannot be sampled efficiently through conventional MD, alternative biased simulation strategies may provide further insights into the primer extension reaction. In addition, our current study is limited by the choice of force field (29) and does not consider the subsequent steps on the reaction pathway such as O3′-P bond formation, expulsion of Mg^2+^ from the reaction center, and the structural dynamics of the leaving group, all of which will require more extensive quantum mechanical studies using QM/MM techniques.

In summary, we have used MD simulations to examine a set of models of the primer extension reaction center in which a Mg^2+^ ion has been placed in alternative coordination geometries. We predict a specific catalytically favorable coordination of the metal ion within the reaction center. This prediction can be tested by thiol-metal substitution experiments, in combination with conventional and time-resolved X-ray crystallographic studies. We suggest that the catalytic Mg^2+^ ion may play multiple roles including facilitating deprotonation of the 3′-hydroxyl and increasing the electrophilicity of the reactive phosphorous. We propose that the catalytic Mg^2+^ also plays an electrostatic role in overcoming repulsion between the negative charge of a deprotonated O3′ and the phosphate being attacked, and plays an organizational role in stabilizing the conformation of the bridged dinucleotide and its position relative to the terminal primer nucleotide.

## AUTHOR INFORMATION

Corresponding Author

^*^

## Author Contributions

The manuscript was written through contributions of all authors. All authors have given approval to the final version of the manuscript.

## Notes

The authors declare no competing financial interest.

## Acknowledgements

Portions of this research were conducted on the O2 High Performance Compute Cluster, supported by the Research Computing Group, at Harvard Medical School.

See https://it.hms.harvard.edu/our-services/research-computing for more information. Remaining simulations were performed using HPC resources on the Midway3 supercomputer at the Research Computing Center at the University of Chicago. We thank Dr. Rafal Szabla for his helpful comments and insights. This work was supported by a grant (290363) from the Simons Foundation to J.W.S. J.W.S. is an Investigator of the Howard Hughes Medical Institute.

## REFERENCES

1. Lohrmann, R., and L.E. Orgel. 1973. Prebiotic Activation Processes. Nature 1973 244:5416. 244:418–420.

2. Inoue, T., G.F. Joyce, K. Grzeskowiak, L.E. Orgel, J.M. Brown, and C.B. Reese. 1984. Template-directed synthesis on the pentanucleotide CpCpGpCpC. Journal of Molecular Biology. 178:669–676.

3. Orgel, L.E. 1992. Molecular replication. Nature 1992 358:6383. 358:203–209.

4. Weimann, B.J., R. Lohrmann, L.E. Orgel, H. Schneider-Bernloehr, and J.E. Sulston. 1968. Template-Directed Synthesis with Adenosine-5′-phosphorimidazolide. Science. 161:387.

5. Inoue, T., and L.E. Orgel. 1982. Oligomerization of (guanosine 5′-phosphor)-2-methylimidazolide on poly(C): An RNA polymerase model. Journal of Molecular Biology. 162:201–217.

6. Jauker, M., H. Griesser, and C. Richert. 2015. Copying of RNA Sequences without Pre-Activation. Angewandte Chemie (International Ed. in English). 54:14559.

7. Vogel, S.R., C. Deck, and C. Richert. 2005. Accelerating chemical replication steps of RNA involving activated ribonucleotides and downstream-binding elements. Chemical Communications. 4922–4924.

8. Vázquez-Salazar, A., G. Tan, A. Stockton, R. Fani, A. Becerra, and A. Lazcano. 2017. Can an Imidazole Be Formed from an Alanyl-Seryl-Glycine Tripeptide under Possible Prebiotic Conditions? Origins of Life and Evolution of Biospheres. 47:345–354.

9. Shen, C., L. Yang, S.L. Miller, and J. Oró. 1987. Prebiotic synthesis of imidazole-4-acetaldehyde and histidine. Origins of life and evolution of the biosphere : the journal of the International Society for the Study of the Origin of Life. 17:295–305.

10. Oró, J., B. Basile, S. Cortes, C. Shen, and T. Yamrom. 1984. The prebiotic synthesis and catalytic role of imidazoles and other condensing agents. Origins of life. 14:237–242.

11. Fahrenbach, A.C., C. Giurgiu, C.P. Tam, L. Li, Y. Hongo, M. Aono, and J.W. Szostak. 2017. Common and Potentially Prebiotic Origin for Precursors of Nucleotide Synthesis and Activation..

12. Ding, D., S.J. Zhang, and J.W. Szostak. 2023. Enhanced nonenzymatic RNA copying with in-situ activation of short oligonucleotides. Nucleic Acids Research. 51:6528.

13. Mariani, A., D.A. Russell, T. Javelle, and J.D. Sutherland. 2018. A Light-Releasable Potentially Prebiotic Nucleotide Activating Agent. Journal of the American Chemical Society. 140:8657–8661.

14. Zhang, S.J., D. Duzdevich, D. Ding, and J.W. Szostak. 2022. Freeze-thaw cycles enable a prebiotically plausible and continuous pathway from nucleotide activation to nonenzymatic RNA copying. Proceedings of the National Academy of Sciences of the United States of America. 119.

15. Li, L., N. Prywes, C.P. Tam, D.K. Oflaherty, V.S. Lelyveld, E.C. Izgu, A. Pal, and J.W. Szostak. 2017. Enhanced nonenzymatic RNA copying with 2-aminoimidazole activated nucleotides. Journal of the American Chemical Society. 139:1810–1813.

16. Zhou, L., S.C. Kim, K.H. Ho, D.K. O’flaherty, C. Giurgiu, T.H. Wright, and J.W. Szostak. 2019. Non-enzymatic primer extension with strand displacement..

17. Walton, T., and J.W. Szostak. 2017. A Kinetic Model of Nonenzymatic RNA Polymerization by Cytidine-5′-phosphoro-2-aminoimidazolide. Biochemistry. 56:5739–5747.

18. Walton, T., and J.W. Szostak. 2016. A Highly Reactive Imidazolium-Bridged Dinucleotide Intermediate in Nonenzymatic RNA Primer Extension. J. Am. Chem. Soc. 138:11996–12002.

19. Walton, T., L. Pazienza, and J.W. Szostak. 2019. Template-Directed Catalysis of a Multistep Reaction Pathway for Nonenzymatic RNA Primer Extension. Biochemistry. 58:755–762.

20. Prywes, N., J.C. Blain, F. Del Frate, and J.W. Szostak. 2016. Nonenzymatic copying of RNA templates containing all four letters is catalyzed by activated oligonucleotides. eLife. 5:e17756.

21. Zhang, W., T. Walton, L. Li, and J.W. Szostak. 2018. Crystallographic observation of nonenzymatic RNA primer extension. eLife. 7:e36422.

22. Joyce, G.F., and L.E. Orgel. 1988. Non-enzymatic template-directed synthesis on RNA random copolymers: Poly(C,A) templates. Journal of Molecular Biology. 202:677–681.

23. Kanavarioti, A., C.F. Bernasconi, D.L. Doodokyan, and D.J. Alberas. 1989. Magnesium ion catalyzed P-N bond hydrolysis in imidazolide-activated nucleotides. Relevance to template-directed synthesis of polynucleotides. Journal of the American Chemical Society. 111:7247–7257.

24. Taifeng, W., and L.E. Orgel. 1992. Nonenzymatic template-directed synthesis on oligodeoxycytidylate sequences in hairpin oligonucleotides. Journal of the American Chemical Society. 114:317–322.

25. Zhang, W., C.P. Tam, L. Zhou, S.S. Oh, J. Wang, and J.W. Szostak. 2018. Structural Rationale for the Enhanced Catalysis of Nonenzymatic RNA Primer Extension by a Downstream Oligonucleotide. J. Am. Chem. Soc. 140:2829–2840.

26. Giurgiu, C., Z. Fang, H.R.M. Aitken, S.C. Kim, L. Pazienza, S. Mittal, and J.W. Szostak. 2021. Structure–Activity Relationships in Nonenzymatic Template-Directed RNA Synthesis. Angewandte Chemie International Edition. 60:22925–22932.

27. Frederiksen, J.K., R. Fong, and J.A. Piccirilli. 2008. Chapter 8. Metal Ions in RNA Catalysis. In: Hud NV, editor. RSC Biomolecular Sciences. Cambridge: Royal Society of Chemistry. pp. 260–306.

28. Giurgiu, C., T.H. Wright, D.K. O’Flaherty, and J.W. Szostak. 2018. A Fluorescent G-Quadruplex Sensor for Chemical RNA Copying. Angewandte Chemie International Edition. 57:9844–9848.

29. Šponer, J., G. Bussi, M. Krepl, P. Banáš, S. Bottaro, R.A. Cunha, A. Gil-Ley, G. Pinamonti, S. Poblete, P. Jurečka, N.G. Walter, and M. Otyepka. 2018. RNA Structural Dynamics As Captured by Molecular Simulations: A Comprehensive Overview. Chem. Rev. 118:4177–4338.

30. Zgarbová, M., M. Otyepka, J. Šponer, F. Lankaš, and P. Jurečka. 2014. Base Pair Fraying in Molecular Dynamics Simulations of DNA and RNA. J. Chem. Theory Comput. 10:3177–3189.

31. Li, L., and J.W. Szostak. 2014. The Free Energy Landscape of Pseudorotation in 3′–5′ and 2′–5′ Linked Nucleic Acids. J. Am. Chem. Soc. 136:2858–2865.

32. Bottaro, S., P.J. Nichols, B. Vögeli, M. Parrinello, and K. Lindorff-Larsen. 2020. Integrating NMR and simulations reveals motions in the UUCG tetraloop. Nucleic Acids Research. 48:5839–5848.

33. Yu, T., and S.-J. Chen. 2018. Hexahydrated Mg2+ Binding and Outer-Shell Dehydration on RNA Surface. Biophysical Journal. 114:1274–1284.

34. Ratnasinghe, B.D., A.M. Salsbury, and J.A. Lemkul. 2020. Ion Binding Properties and Dynamics of the bcl-2 G-Quadruplex Using a Polarizable Force Field. J. Chem. Inf. Model. 60:6476–6488.

35. Zhang, S., D.R. Stevens, P. Goyal, J.L. Bingaman, P.C. Bevilacqua, and S. Hammes-Schiffer. 2016. Assessing the Potential Effects of Active Site Mg2+ Ions in the glmS Ribozyme–Cofactor Complex. J. Phys. Chem. Lett. 7:3984–3988.

36. Ganguly, A., B.P. Weissman, T.J. Giese, N.-S. Li, S. Hoshika, S. Rao, S.A. Benner, J.A. Piccirilli, and D.M. York. 2020. Confluence of theory and experiment reveals the catalytic mechanism of the Varkud satellite ribozyme. Nat. Chem. 12:193–201.

37. Ucisik, M.N., P.C. Bevilacqua, and S. Hammes-Schiffer. 2016. Molecular Dynamics Study of Twister Ribozyme: Role of Mg2+ Ions and the Hydrogen-Bonding Network in the Active Site. Biochemistry. 55:3834–3846.

38. Macke, T.J., and D.A. Case. 1998. Modeling Unusual Nucleic Acid Structures. ACS Symposium Series. 682:379–393.

39. Humphrey, W., A. Dalke, and K. Schulten. 1996. VMD: visual molecular dynamics. J Mol Graph. 14:33–38, 27–28.

40. MacKerell, A.D., D. Bashford, M. Bellott, R.L. Dunbrack, J.D. Evanseck, M.J. Field, S. Fischer, J. Gao, H. Guo, S. Ha, D. Joseph-McCarthy, L. Kuchnir, K. Kuczera, F.T.K. Lau, C. Mattos, S. Michnick, T. Ngo, D.T. Nguyen, B. Prodhom, W.E. Reiher, B. Roux, M. Schlenkrich, J.C. Smith, R. Stote, J. Straub, M. Watanabe, J. Wiórkiewicz-Kuczera, D. Yin, and M. Karplus. 1998. All-Atom Empirical Potential for Molecular Modeling and Dynamics Studies of Proteins. J. Phys. Chem. B. 102:3586–3616.

41. Denning, E.J., U.D. Priyakumar, L. Nilsson, A.D. MacKerell, and Jr. 2011. Impact of 2′-hydroxyl sampling on the conformational properties of RNA: Update of the CHARMM all-atom additive force field for RNA. Journal of computational chemistry. 32:1929.

42. Vanommeslaeghe, K., E. Hatcher, C. Acharya, S. Kundu, S. Zhong, J. Shim, E. Darian, O. Guvench, P. Lopes, I. Vorobyov, and A.D. Mackerell. 2010. CHARMM general force field: A force field for drug-like molecules compatible with the CHARMM all-atom additive biological force fields. J Comput Chem. 31:671–690.

43. Bignon, E., and A. Monari. 2022. Modeling the Enzymatic Mechanism of the SARS-CoV-2 RNA-Dependent RNA Polymerase by DFT/MM-MD: An Unusual Active Site Leading to High Replication Rates. J. Chem. Inf. Model. 62:4261–4269.

44. Phillips, J.C., D.J. Hardy, J.D.C. Maia, J.E. Stone, J.V. Ribeiro, R.C. Bernardi, R. Buch, G. Fiorin, J. Hénin, W. Jiang, R. McGreevy, M.C.R. Melo, B.K. Radak, R.D. Skeel, A. Singharoy, Y. Wang, B. Roux, A. Aksimentiev, Z. Luthey-Schulten, L.V. Kalé, K. Schulten, C. Chipot, and E. Tajkhorshid. 2020. Scalable molecular dynamics on CPU and GPU architectures with NAMD. J. Chem. Phys. 153:044130.

45. Fiorin, G., M.L. Klein, and J. Hénin. 2013. Using collective variables to drive molecular dynamics simulations. Molecular Physics. 111:3345–3362.

46. Martínez, L., R. Andrade, E.G. Birgin, and J.M. Martínez. 2009. PACKMOL: a package for building initial configurations for molecular dynamics simulations. J Comput Chem. 30:2157–2164.

47. Roe, D.R., and T.E. Cheatham. 2013. PTRAJ and CPPTRAJ: Software for Processing and Analysis of Molecular Dynamics Trajectory Data. J. Chem. Theory Comput. 9:3084–3095.

48. McGibbon, R.T., K.A. Beauchamp, M.P. Harrigan, C. Klein, J.M. Swails, C.X. Hernández, C.R. Schwantes, L.-P. Wang, T.J. Lane, and V.S. Pande. 2015. MDTraj: A Modern Open Library for the Analysis of Molecular Dynamics Trajectories. Biophysical Journal. 109:1528–1532.

49. Altona, C., and M. Sundaralingam. 1972. Conformational analysis of the sugar ring in nucleosides and nucleotides. New description using the concept of pseudorotation. J. Am. Chem. Soc. 94:8205–8212.

50. Zgarbová, M., M. Otyepka, J. Šponer, A. Mládek, P. Banáš, T.E. Cheatham, and P. Jurečka. 2011. Refinement of the Cornell et al. Nucleic acids force field based on reference quantum chemical calculations of glycosidic torsion profiles. Journal of Chemical Theory and Computation. 7:2886–2902.

51. Jo, S., T. Kim, V.G. Iyer, and W. Im. 2008. CHARMM-GUI: A web-based graphical user interface for CHARMM. Journal of Computational Chemistry. 29:1859–1865.

52. Lee, J., M. Hitzenberger, M. Rieger, N.R. Kern, M. Zacharias, and W. Im. 2020. CHARMM-GUI supports the Amber force fields. The Journal of Chemical Physics. 153:035103.

53. Lemkul, J.A., and A.D. Mackerell. 2016. Balancing the Interactions of Mg2+ in Aqueous Solution and with Nucleic Acid Moieties For a Polarizable Force Field Based on the Classical Drude Oscillator Model. The journal of physical chemistry. B. 120:11436.

54. Melo, M.C.R., R.C. Bernardi, T. Rudack, M. Scheurer, C. Riplinger, J.C. Phillips, J.D.C. Maia, G.B. Rocha, J.V. Ribeiro, J.E. Stone, F. Neese, K. Schulten, and Z. Luthey-Schulten. 2018. NAMD goes quantum: An integrative suite for QM/MM simulations. Nature methods. 15:351.

55. Neese, F., F. Wennmohs, U. Becker, and C. Riplinger. 2020. The ORCA quantum chemistry program package. The Journal of Chemical Physics. 152:224108.

56. Sure, R., and S. Grimme. 2013. Corrected small basis set Hartree-Fock method for large systems. Journal of Computational Chemistry. 34:1672–1685.

57. Zhang, W., C.P. Tam, T. Walton, A.C. Fahrenbach, G. Birrane, and J.W. Szostak. 2017. Insight into the mechanism of nonenzymatic RNA primer extension from the structure of an RNA-GpppG complex. PNAS. 114:7659–7664.

58. Wu, T., and L.E. Orgel. 1992. Nonenzymic template-directed synthesis on oligodeoxycytidylate sequences in hairpin oligonucleotides. J. Am. Chem. Soc. 114:317–322.

59. Kozlov, I.A., P.K. Politis, A. Van Aerschot, R. Busson, P. Herdewijn, and L.E. Orgel. 1999. Nonenzymatic Synthesis of RNA and DNA Oligomers on Hexitol Nucleic Acid Templates: The Importance of the A Structure. J. Am. Chem. Soc. 121:2653–2656.

60. Zheng, H., I.G. Shabalin, K.B. Handing, J.M. Bujnicki, and W. Minor. 2015. Magnesium-binding architectures in RNA crystal structures: validation, binding preferences, classification and motif detection. Nucleic Acids Research. 43:3789–3801.

61. Sawle, L., and K. Ghosh. 2016. Convergence of Molecular Dynamics Simulation of Protein Native States: Feasibility vs Self-Consistency Dilemma. Journal of Chemical Theory and Computation. 12:861–869.

62. Ding, D., L. Zhou, C. Giurgiu, and J.W. Szostak. 2022. Kinetic explanations for the sequence biases observed in the nonenzymatic copying of RNA templates. Nucleic Acids Research. 50:35–45.

63. Giurgiu, C., L. Li, D.K. O’Flaherty, C.P. Tam, and J.W. Szostak. 2017. A Mechanistic Explanation for the Regioselectivity of Nonenzymatic RNA Primer Extension. J. Am. Chem. Soc. 139:16741–16747.

64. Steffen, F.D., M. Khier, D. Kowerko, R.A. Cunha, R. Börner, and R.K.O. Sigel. 2020. Metal ions and sugar puckering balance single-molecule kinetic heterogeneity in RNA and DNA tertiary contacts. Nature Communications 2020 11:1. 11:1–11.

65. Duzdevich, D., C.E. Carr, D. Ding, S.J. Zhang, T.S. Walton, and J.W. Szostak. 2021. Competition between bridged dinucleotides and activated mononucleotides determines the error frequency of nonenzymatic RNA primer extension. Nucleic Acids Research. 49:3681–3691.

66. Thaplyal, P., A. Ganguly, S. Hammes-Schiffer, and P.C. Bevilacqua. 2015. Inverse Thio Effects in the Hepatitis Delta Virus Ribozyme Reveal that the Reaction Pathway Is Controlled by Metal Ion Charge Density. Biochemistry. 54:2160–2175.

67. Kim, S.C., L. Zhou, W. Zhang, D.K. O’Flaherty, V. Rondo-Brovetto, and J.W. Szostak. 2020. A Model for the Emergence of RNA from a Prebiotically Plausible Mixture of Ribonucleotides, Arabinonucleotides, and 2′-Deoxynucleotides. J. Am. Chem. Soc. 142:2317–2326.

68. Zhang, W., S.C. Kim, C.P. Tam, V.S. Lelyveld, S. Bala, J.C. Chaput, and J.W. Szostak. 2021. Structural interpretation of the effects of threo-nucleotides on nonenzymatic template-directed polymerization. Nucleic Acids Research. 49:646–656.

69. Kim, S.C., D.K. O’Flaherty, C. Giurgiu, L. Zhou, and J.W. Szostak. 2021. The Emergence of RNA from the Heterogeneous Products of Prebiotic Nucleotide Synthesis. J. Am. Chem. Soc. 143:3267–3279.

