## Supplementary Information for "Simulations predict preferred Mg^2+^ coordination in a nonenzymatic primer extension reaction center"

### Supplemental Figures

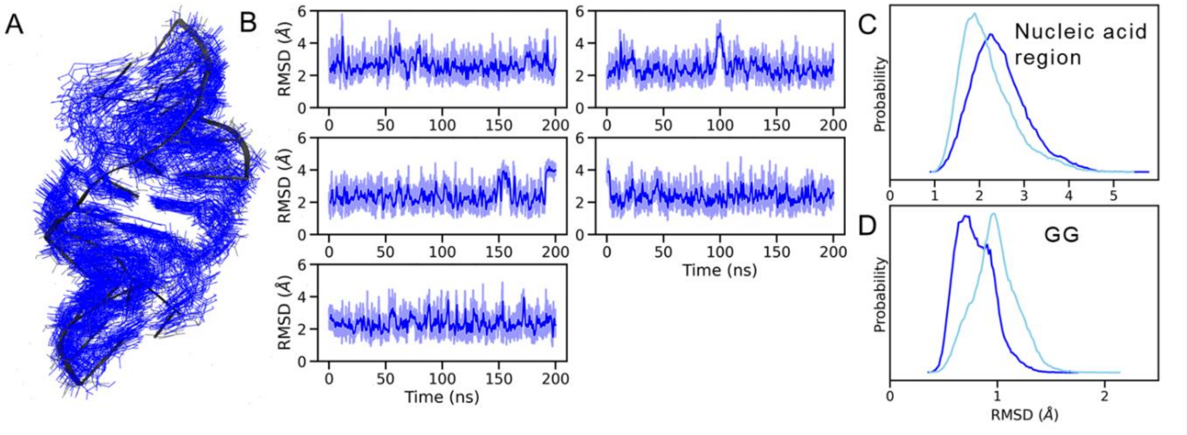


***Figure S1.*** **RMSD of the duplex and reaction center suggest increased dynamics with a deprotonated 3′-OH.** (A) Overlay of simulation snapshots over a 500 ns simulation trajectory of the complex with primer 3′-OH w/o $Mg^{2+}$. The first frame of the simulation can be seen partly as a dark gray ribbon representation. (B) Time series of root mean-squared deviation (RMSD) for five simulation replicates. Comparing RMSD probability distribution for the (C) nucleic acid region and (D) GG bridged dinucleotide for the simulation systems, 3′-OH w/o $Mg^{2+}$ (dark blue) and 3′$O^-$ w/o $Mg^{2+}$ (light blue).

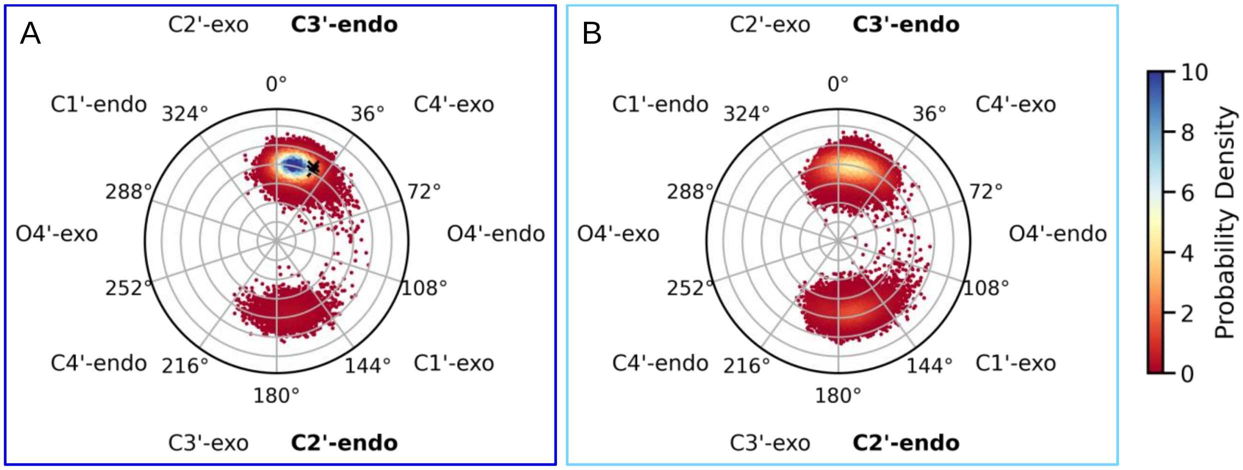


***Figure S2.*** **Terminal primer sugar puckering in the absence of bound $Mg^{2+}$ and a major groove facing 2AI.** Circular histogram of pseudorotation angles and amplitudes of the terminal primer nucleotide sugar for the (A) 3′-OH w/o $Mg^{2+}$ (dark blue) and (B) 3′$O^-$ w/o $Mg^{2+}$ (light blue) simulation systems. The phase angles are based on the Altona-Sundaralingam[1] definition and are assigned to the puckering modes in multiples of 36°. Sugar pucker conformations of the primer nucleotide from the crystal structure, denoted with × markers in the circular plot show the sugars are in the C3′-endo conformation.

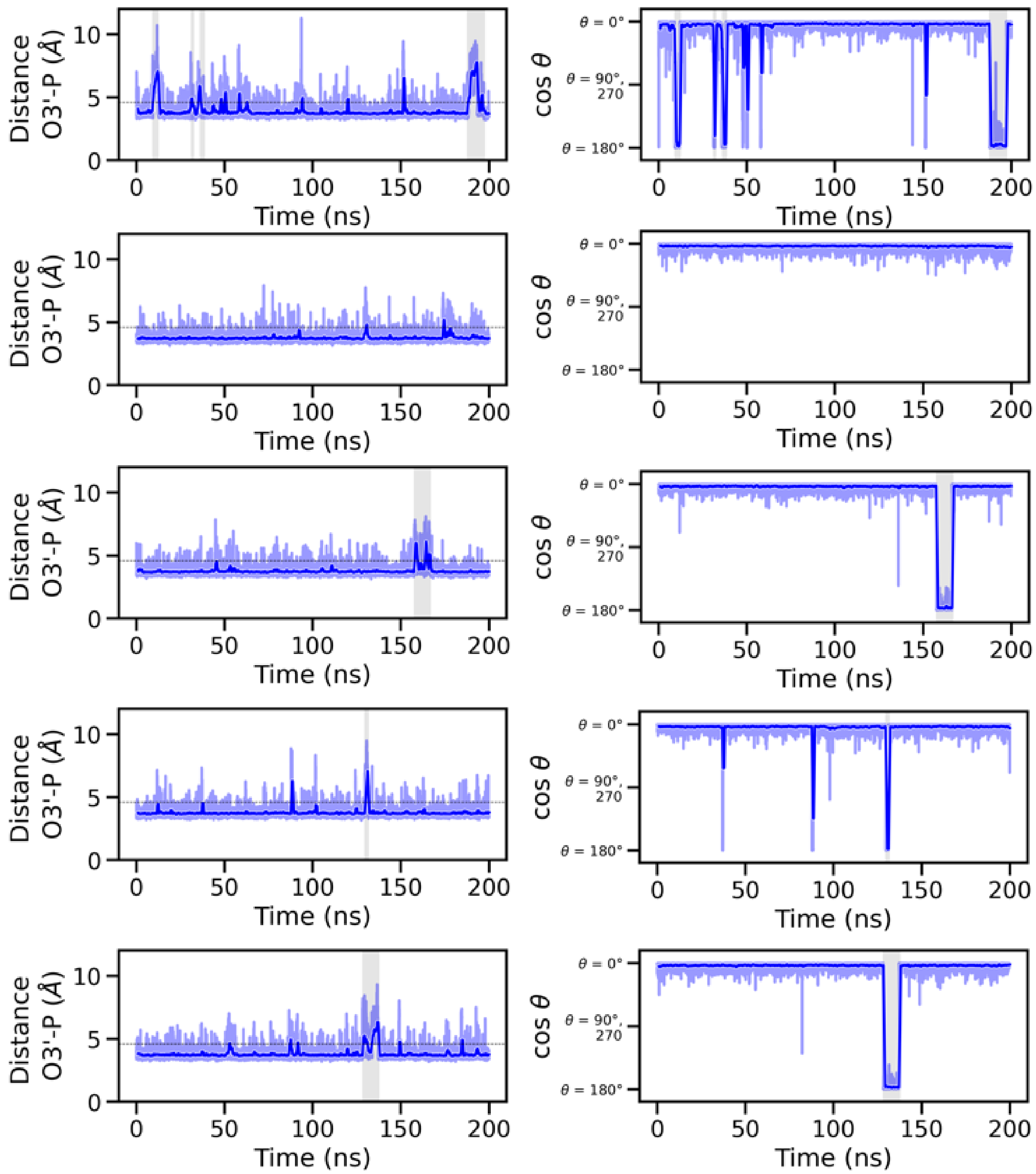


***Figure S3.*** **Time series of the distance O3′ (primer)-P (bridged dinucleotide) and primer nucleotide sugar pucker angle for the 3′-OH w/o $Mg^{2+}$ simulation system for all five replicates.** The horizontal dashed lines indicate the crystal structure value. Intervals with larger distance values and C2′-endo sugar conformations are highlighted in gray. For all time series plots, dark traces show the data averaged over a 1 ns window, while the lighter envelope shows the full range of the data recorded at 10 ps time steps in our simulations.

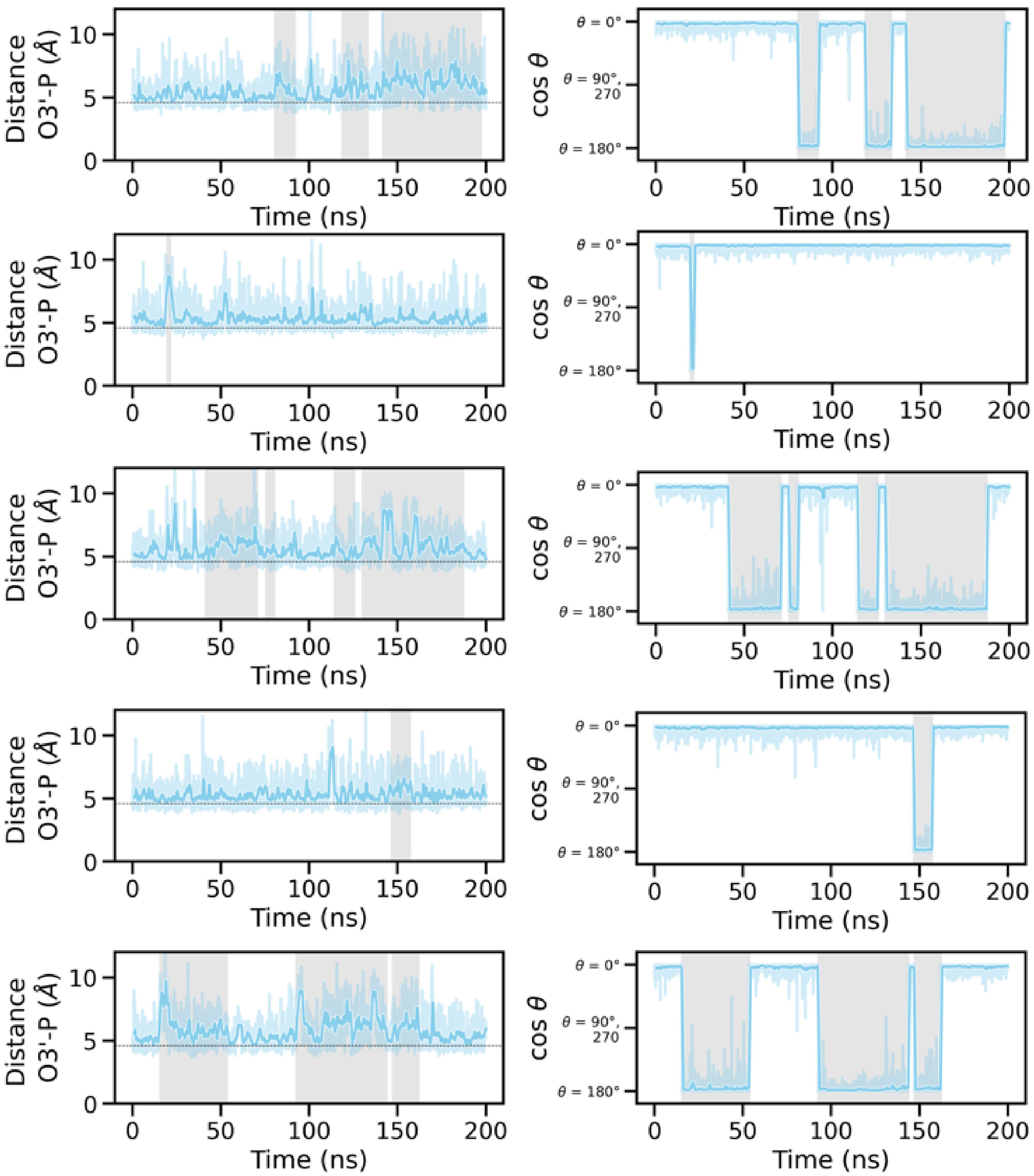


***Figure S4.*** **Time series of the distance O3′ (primer)-P (bridged dinucleotide) and primer nucleotide sugar pucker angle for the 3′O- w/o $Mg^{2+}$ simulation system for all five replicates.** The horizontal dashed lines indicate the crystal structure value. Intervals with larger distance values and C2′-endo sugar conformations are highlighted in gray. For all time series plots, dark traces show the data averaged over a 1 ns window, while the lighter envelope shows the full range of the data recorded at 10 ps time steps in our simulations.

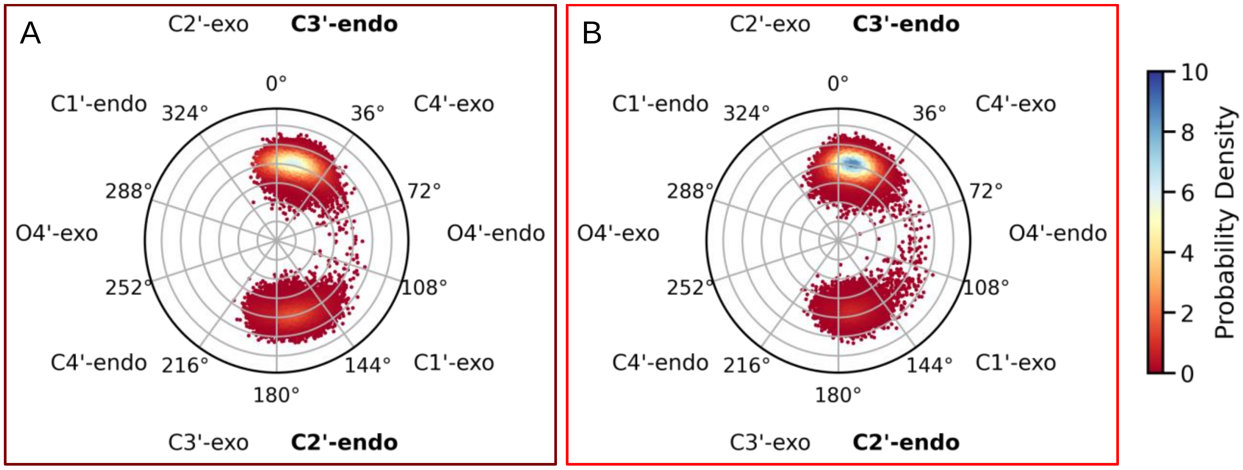


***Figure S5.*** **Terminal primer sugar puckering in the absence of bound $Mg^{2+}$ and a minor groove facing 2AI.** Circular histogram of pseudorotation angles of the terminal primer nucleotide sugar for the (A) 3′-OH w/o $Mg^{2+}$ (brown) and (B) 3′O- w/o $Mg^{2+}$ (red) simulation systems where 2-$NH_2$-Im orientation is minor groove-facing. The phase angles are based on the Altona-Sundaralingam[1] definition and are assigned to the puckering modes in multiples of 36°.

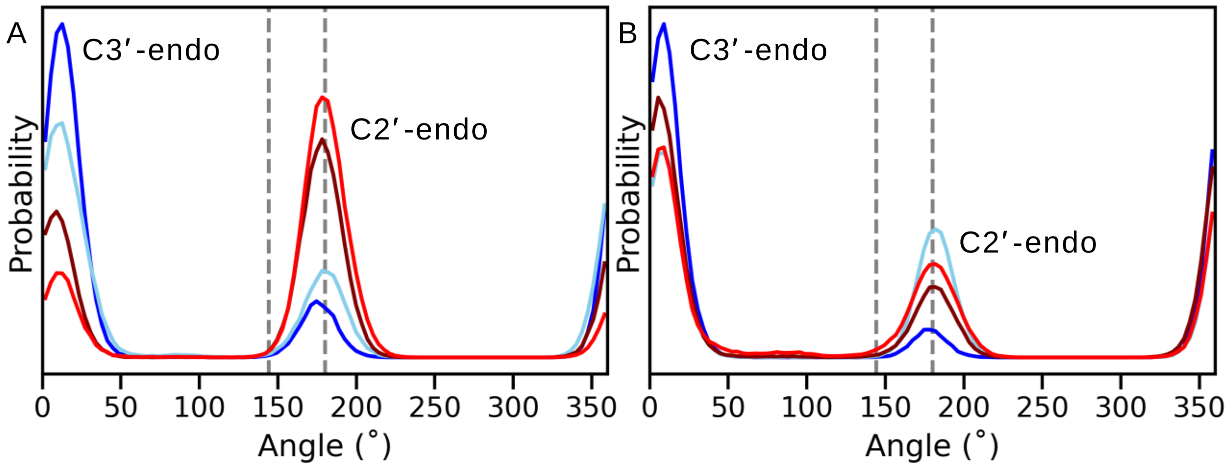


***Figure S6.*** **Probability distribution of the pseudorotation angle of the bridged dinucleotide sugars in the absence of bound $Mg^{2+}$.** (A) G1 and (B) G2 positions in the major groove-facing conformations are shown for 3′-OH w/o $Mg^{2+}$ (dark blue) and 3′O- w/o $Mg^{2+}$ (light blue) and minor groove-facing conformations for 3′-OH w/o $Mg^{2+}$ (brown) and 3′O- w/o $Mg^{2+}$ (red). Dashed lines mark the region for C2′-endo sugar pucker.

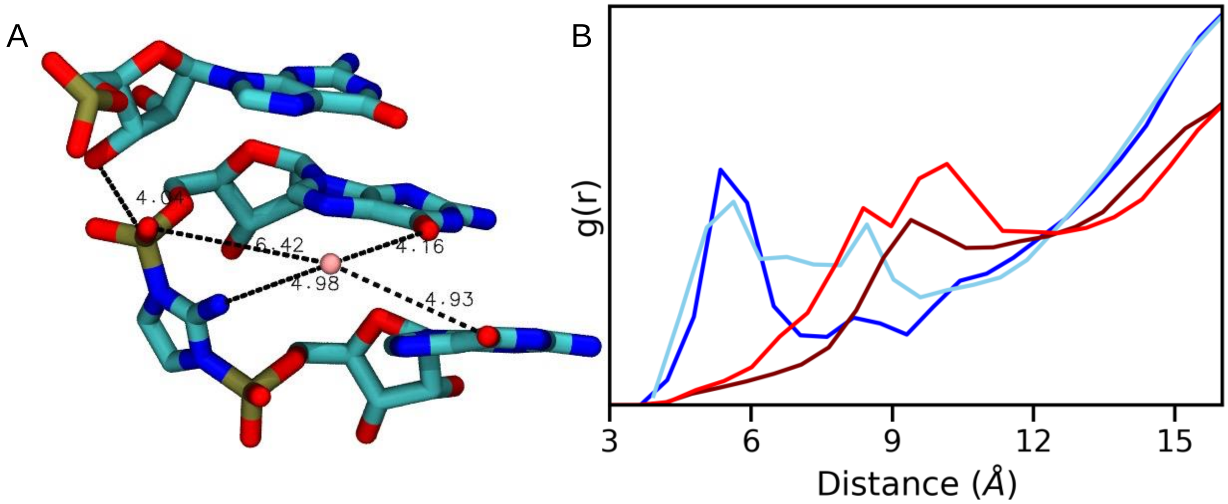


***Figure S7.*** **Water mediated contacts between the bridging imidazolium and $Mg^{2+}$.** (A) Distances of $Mg^{2+}$ ion from various nucleobase atoms and 2-$NH_2$-Im atom of the bridged nucleotide. (B) Radial distribution function of the distance between 2-$NH_2$-Im and $Mg^{2+}$ ions in the MD simulation box. Colors correspond to systems as in Fig. S6.

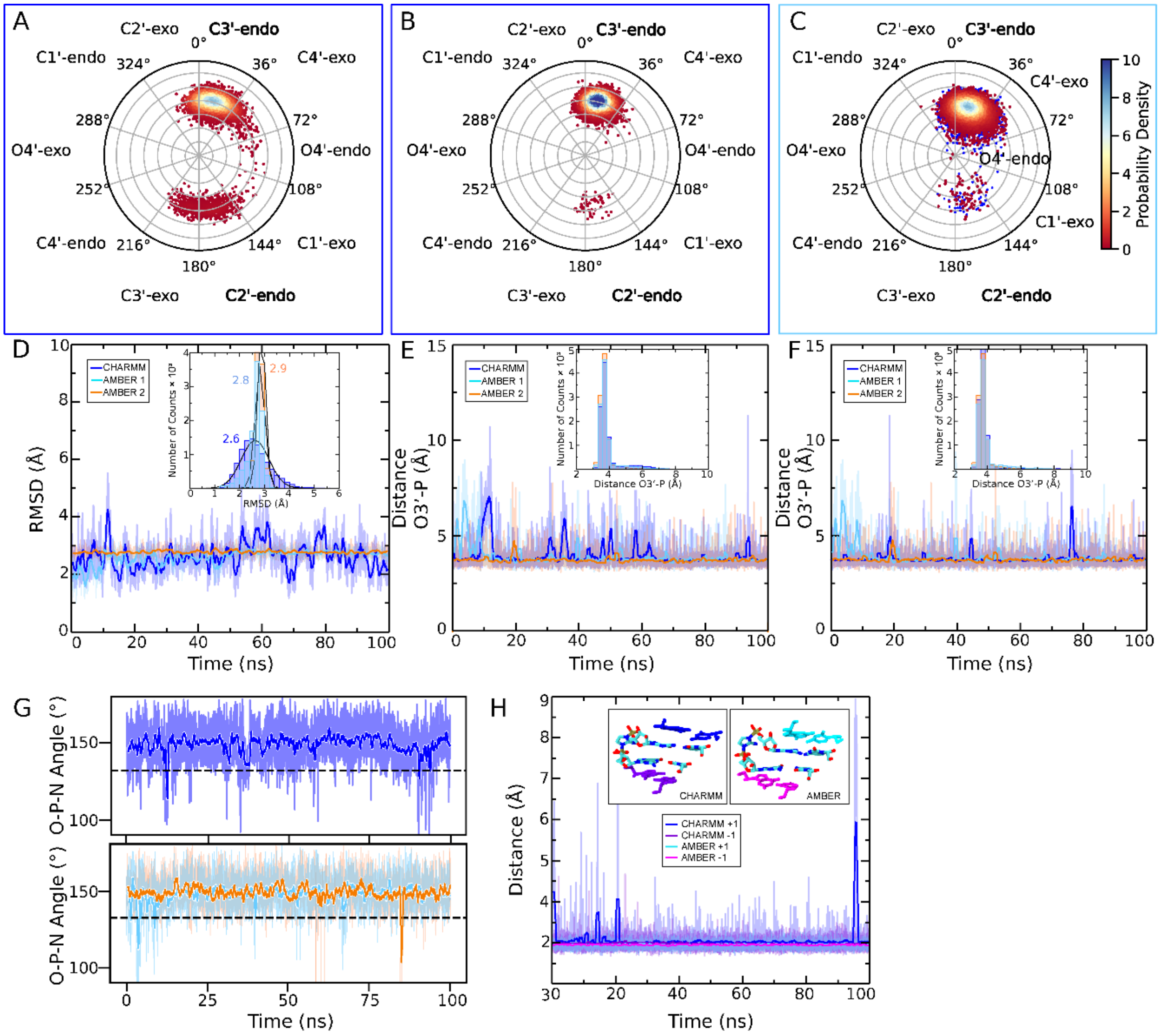


***Figure S8.*** **Comparison of CHARMM and AMBER 3′-OH w/o $Mg^{2+}$ simulations where 2-$NH_2$-Im orientation is major groove-facing.** Terminal primer sugar puckering for (A) 0-100 ns CHARMM (dark blue), (B) 75-175 ns CHARMM (dark blue), and (C) AMBER (light blue) simulation systems. In (C), data points from the first 100 ns are shown on top, while data points from the second 100 ns are shown underneath in blue. The phase angles are based on the Altona-Sundaralingam[1] definition and are assigned to the puckering modes in multiples of 36°. (D) RMSD values of the 0-100 ns CHARMM and two 100 ns AMBER sub-trajectories. The inset shows a probability distribution for each, with mean values indicated in the corresponding color, obtained from a normal fit. Time series of the O3′ (primer)-P (bridged dinucleotide) distance for the 0-100 ns CHARMM and two 100 ns AMBER sub-trajectories (E), and the 75-175 ns CHARMM and two 100 ns AMBER sub-trajectories (F). The insets show a probability distribution for each. (G) The angle of attack during the 0-100 ns CHARMM (top) and two 100 ns AMBER (bottom) sub-trajectories, measured between O3′-P-N atoms. (H) Time series of the distances d(N1,N3) between the G-C base pairs flanking the bridged dinucleotide in the 0-100 ns CHARMM and AMBER simulations. The inset indicates which color traces correspond to the upstream and downstream positions for each simulation.

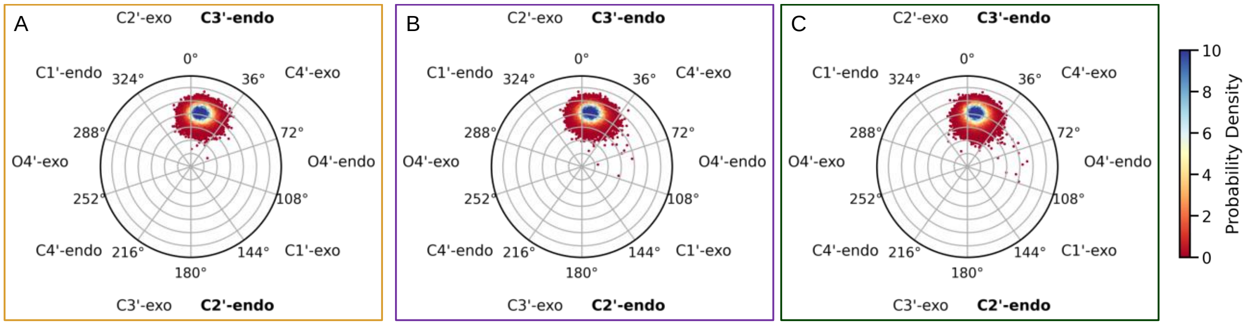


***Figure S9.*** **Terminal primer sugar puckering with bound $Mg^{2+}$.** Circular histogram of pseudorotation angles of the terminal primer nucleotide sugar for the (A) 3′O- w/ $Mg^{2+}$@$S_P$, 2-$NH_2$-Im:$R_P$ (yellow), (B) 3′O- w/ $Mg^{2+}$@$R_P$, 2-$NH_2$-Im:$R_P$ (purple), and (C) 3′O- w/ $Mg^{2+}$@$R_P$, 2-$NH_2$-Im:$S_P$ (green) simulation systems. The phase angles are based on the Altona-Sundaralingam[1] definition and are assigned to the puckering modes in multiples of 36°.

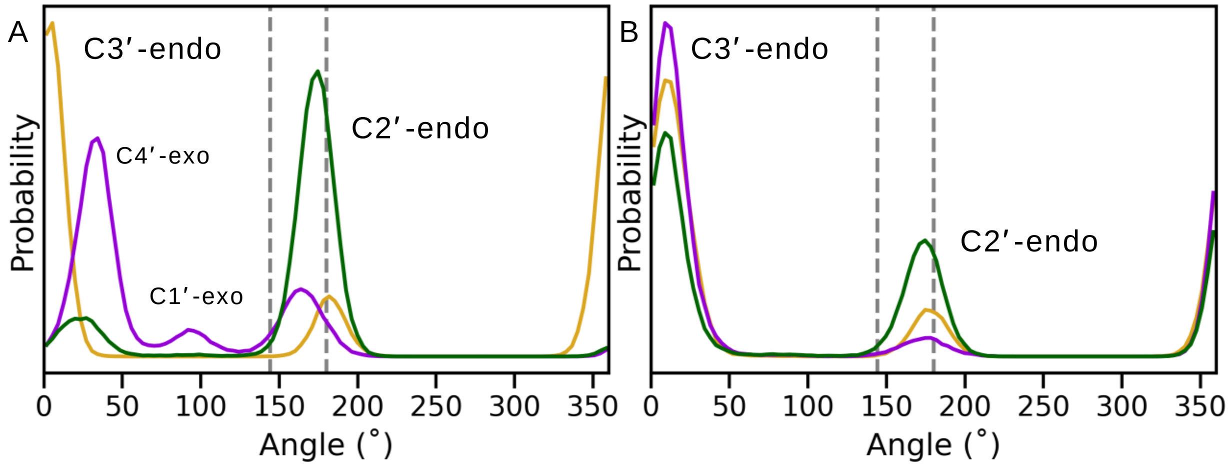


***Figure S10.*** **Probability distribution of the pseudorotation angle of the bridged dinucleotide sugars with bound $Mg^{2+}$.** Probability distribution of the pseudorotation angle of the bridged dinucleotide sugars in the (A) G1 and (B) G2 positions for the simulation ensembles with $Mg^{2+}$ in the reaction center: 3′O- w/ $Mg^{2+}$@$S_P$, 2-$NH_2$-Im:$R_P$ (yellow), 3′O- w/ $Mg^{2+}$@$R_P$, 2-$NH_2$-Im:$R_P$ (purple) and 3′O- w/ $Mg^{2+}$@$R_P$, 2-$NH_2$-Im:$S_P$ (green). Dashed lines mark the region for C2′-endo sugar pucker.

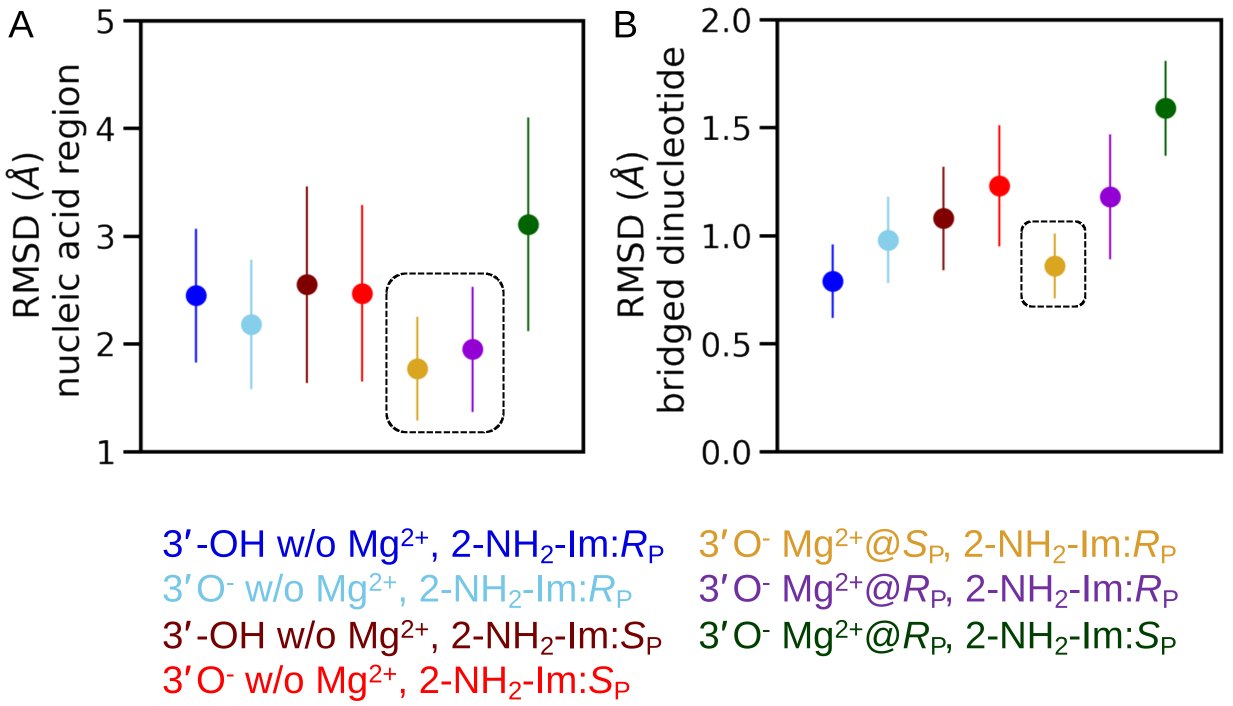


***Figure S11.*** **A bridging $Mg^{2+}$ in the reaction center stabilizes the primer extension complex.** Comparing mean ± standard deviation RMSD of (A) nucleic acid region and (B) bridged dinucleotide heavy atoms among the seven simulation ensembles discussed in this work. Systems highlighted in the main text are outlined.

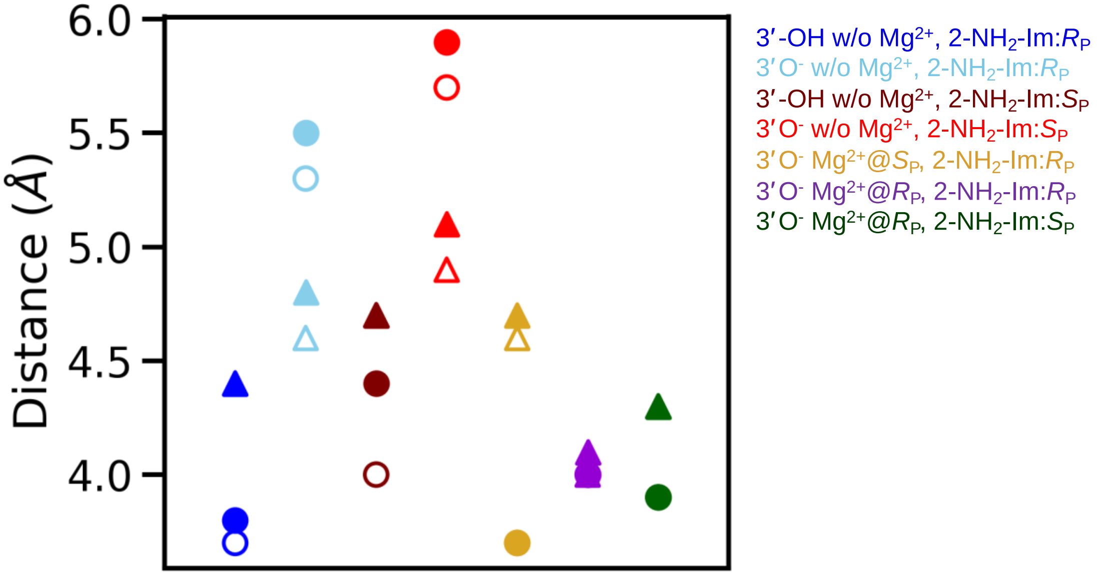


***Figure S12.*** **Distance between O2′-P (△or▲) and O3′-P (○ or ●) for the seven simulation ensembles discussed in this work.** Open and closed markers show mean and median values respectively. Light blue and red highlighted simulation systems show a reversed trend where the average distance(O2′-P) < distance(O3′-P) whereas distance(O2′-P) ≈ distance(O3′-P) for the purple highlighted simulation system.

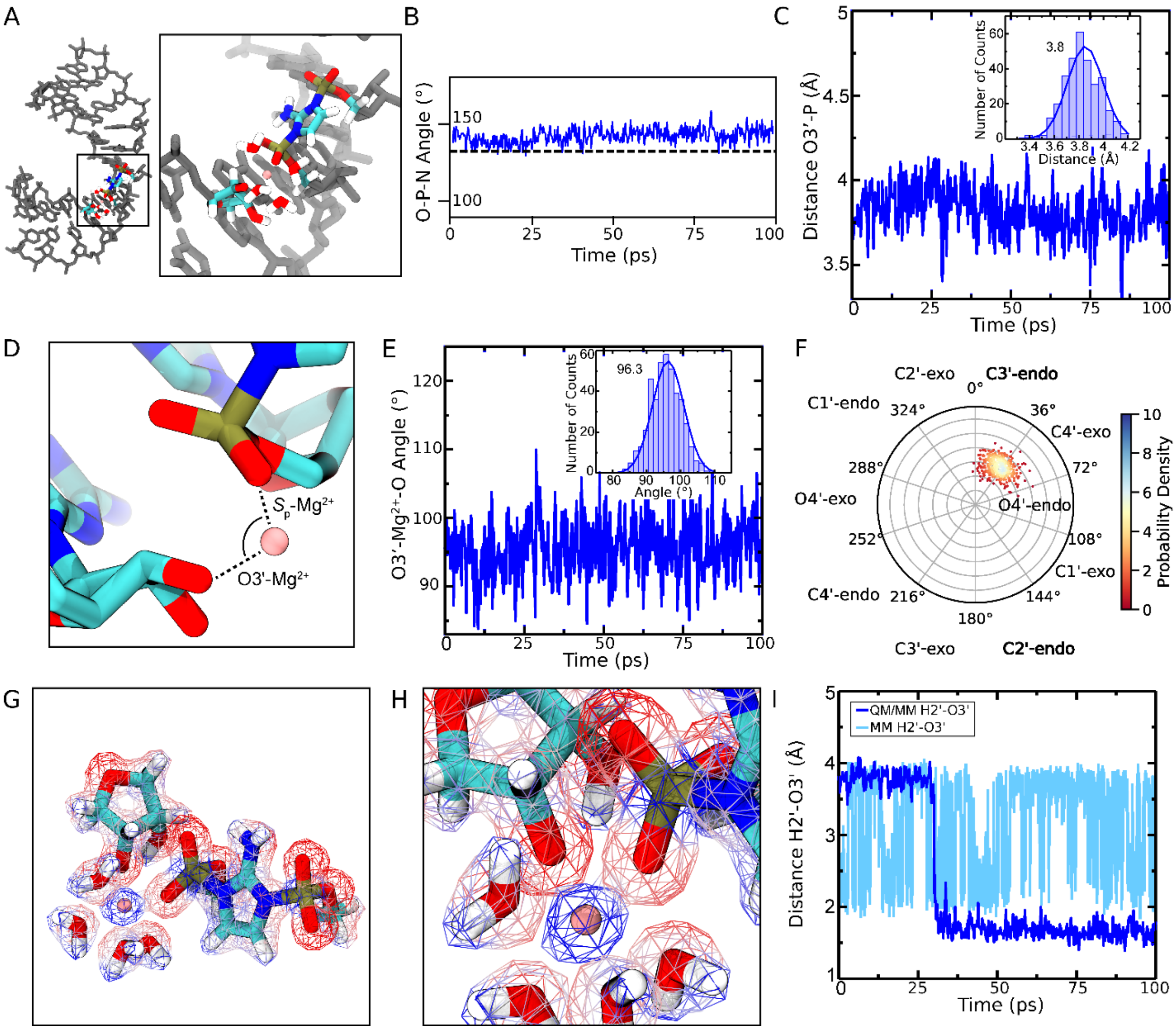


***Figure S13*. QM/MM simulations of the 3′O⁻ w/ $Mg^{2+}$@$S_P$, 2-$NH_2$-Im:$R_P$ system.** (A) Molecular figure of the simulation system in licorice representation showing the portion of the duplex modeled in MM (grey) and the 49 atoms modeled using QM (colored by atom type). (B) The angle of attack during the QM/MM simulation, measured between O3′-P-N atoms, which agrees well with the CHARMM simulations (Figure 5B). (C) Time series of the O3′ (primer)-P (bridged dinucleotide) distance during the QM/MM simulation. The inset shows a probability distribution with the mean value indicated, obtained from a normal fit. This distance agrees well with the average distance from the CHARMM simulations (Figure 4D). (D) Molecular figure showing the O3′-$Mg^{2+}$-O($S_p$) angle measured during the QM/MM simulation (E) Time course of the angle measured in (D). The inset shows a probability distribution with the mean value indicated, obtained from a normal fit. This angle agrees well with the average distance from the CHARMM simulations (Figure S17). (F) Circular histogram of pseudorotation angles of the terminal primer nucleotide sugar during the QM/MM simulation. The phase angles are based on the Altona-Sundaralingam[1] definition and are assigned to the puckering modes in multiples of 36°. (G) A molecular figure of the QM system with calculated SCF electron density colored by the electrostatic potential with positive charge in blue and negative charge in red. (H) A zoomed in view of (G), showing the polarization on the H2′ and O3′. (I) Time series of the H2′ and O3′ distance for the MM and QM/MM simulations, showing the tighter interaction in QM/MM due to polarization of the H2′. The time indicated corresponds to the QM/MM simulations, while the trace corresponding to the MM simulation goes to 3 ns.

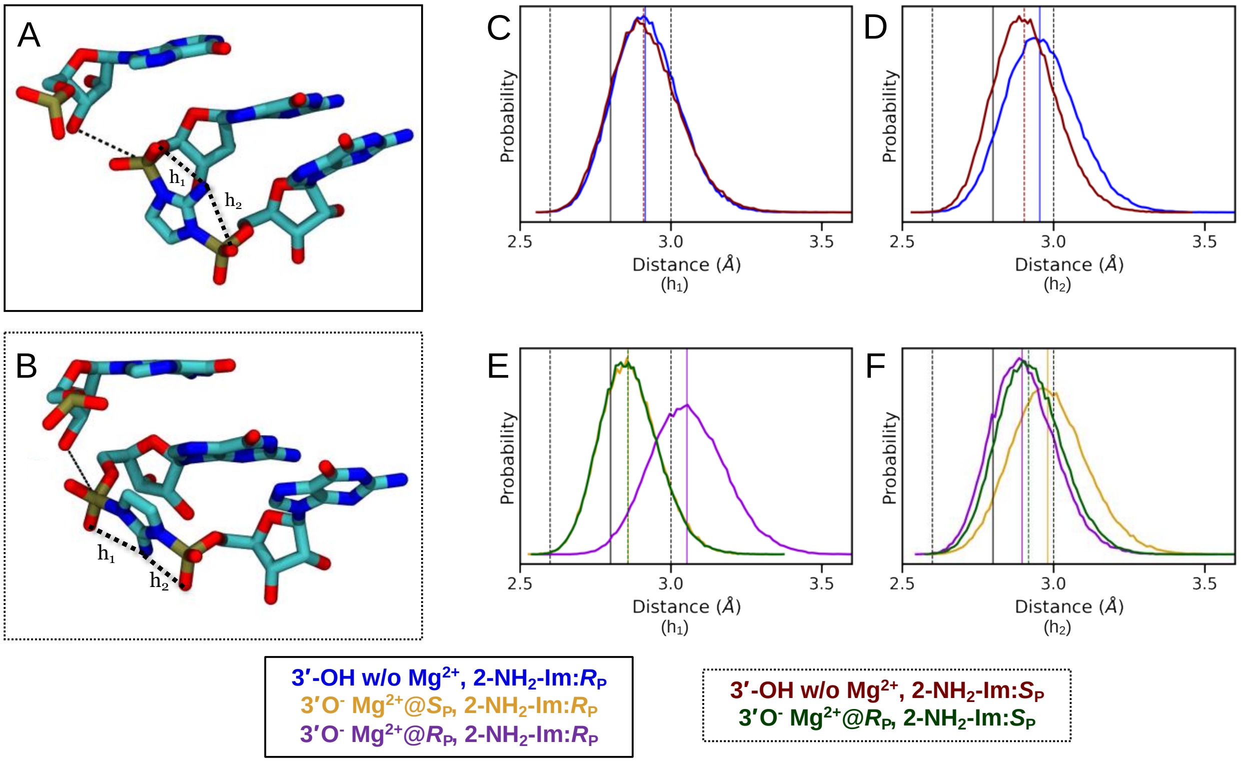


***Figure S14.*** **Hydrogen bonds $h_1$ and $h_2$ between 2-$NH_2$-Im and non-bridging oxygens on the bridged dinucleotide suggest a preferred preorganization geometry.** (A) The major groove-facing and (B) minor groove-facing orientation of the bridged dinucleotide are shown. (C-F) Probability distribution of the hydrogen bond distances $h_1$ and $h_2$ for the various simulation systems. Median distance values from simulation with major and minor groove-facing orientations of bridged dinucleotide are shown in solid and dashed lines, respectively. Lines in black show hydrogen bond distances observed in the crystal structure.

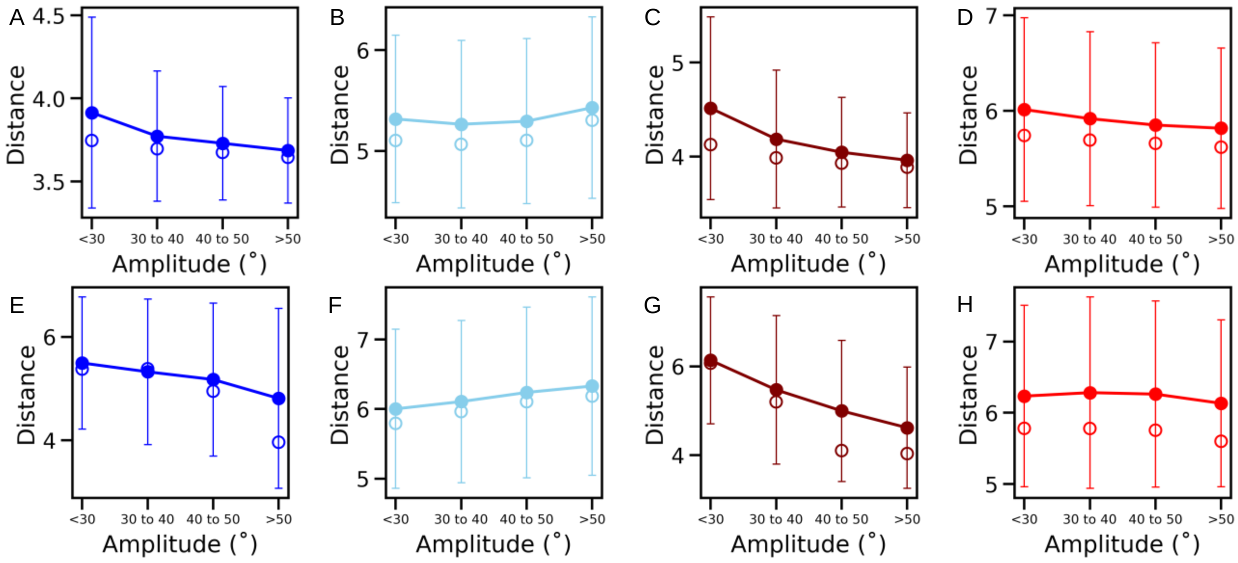


***Figure S15.*** **O3′-P distance as a function of puckering amplitude without bound $Mg^{2+}$.** (A-D) Mean (●) and median (○) of the O3′-P distance distributions when the terminal primer sugar is in the C3′-endo conformation with amplitudes <30°, 30-40°, 40-50°, and >50°. (E-H) Mean (●) and median (○) of the O3′-P distance distributions when the terminal primer sugar is in the C2′-endo conformation with amplitudes <30°, 30-40°, 40-50°, and >50°. Simulation ensembles include major groove-facing 3′-OH (dark blue) and $3'O^-$ (light blue), and minor groove-facing 3′-OH (brown) and $3'O^-$ (red). Standard deviation is shown as colored bars.

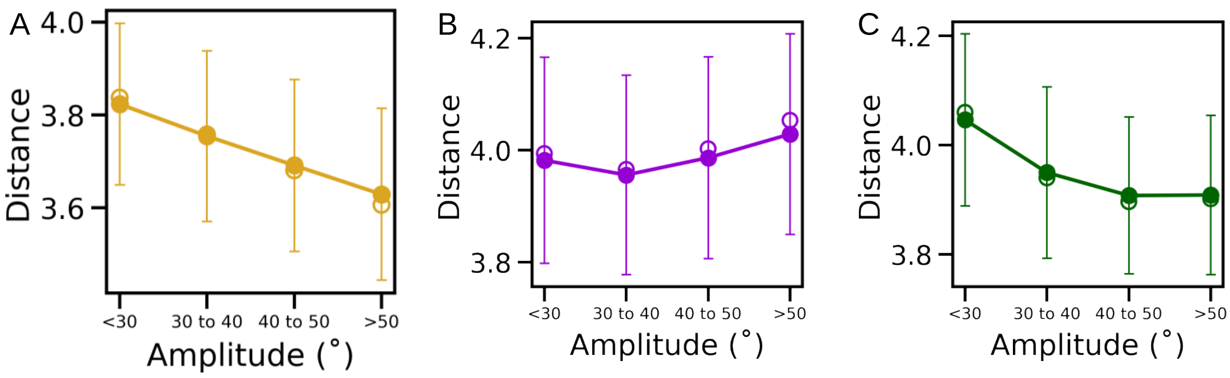


***Figure S16.*** **O3′-P distance as a function of puckering amplitude with bound $Mg^{2+}$.** Mean (●) and median (○) of the O3′-P distance distributions when the terminal primer sugar is in the C3′-endo conformation with amplitudes <30°, 30-40°, 40-50°, and >50°. Simulation ensembles include (A) $3'O^-$ w/ $Mg^{2+}$@$S_P$, 2-$NH_2$-Im:$R_P$ (yellow), (B) $3'O^-$ w/ $Mg^{2+}$@$R_P$, 2-$NH_2$-Im:$R_P$ (purple) and (C) $3'O^-$ w/ $Mg^{2+}$@$R_P$, 2-$NH_2$-Im:$S_P$ (green). Simulation frames in C2′-endo are not observed. Standard deviation is shown as colored bars.

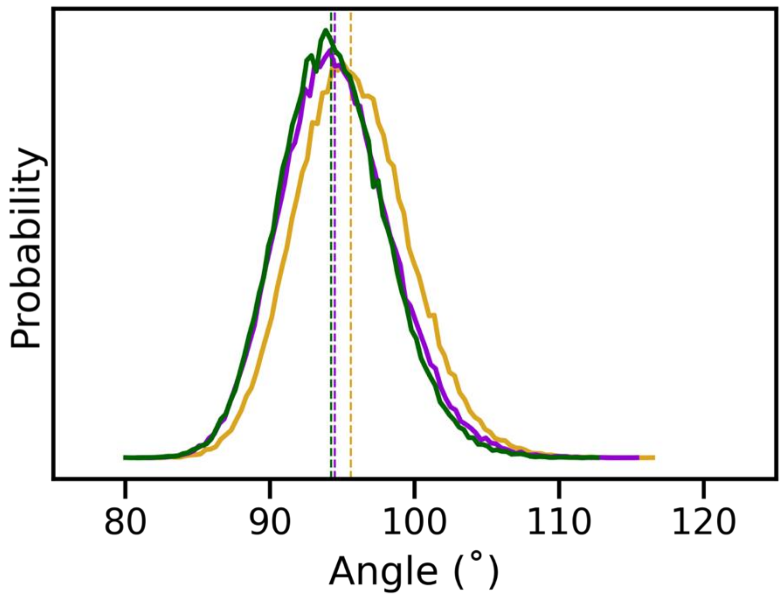


***Figure S17.*** **Probability distribution of the $Mg^{2+}$ coordination angle.** Angles measured between the O3′-$Mg^{2+}$-O ($R_P$ or $S_P$) atoms in three simulation ensembles: 3′$O^-$ w/ $Mg^{2+}$@$S_P$, 2-$NH_2$-Im:$R_P$ (yellow), 3′$O^-$ w/ $Mg^{2+}$@$R_P$, 2-$NH_2$-Im:$R_P$ (purple) and 3′$O^-$ w/ $Mg^{2+}$@$R_P$, 2-$NH_2$-Im:$S_P$ (green). Dashed lines show median values for the distributions.

### REFERENCES

(1) Altona, C.; Sundaralingam, M. Conformational Analysis of the Sugar Ring in Nucleosides and Nucleotides. New Description Using the Concept of Pseudorotation. *J. Am. Chem. Soc.* **1972**, *94* (23), 8205–8212. https://doi.org/10.1021/ja00778a043.
